# Early pre-neural serotonin modulates balance of late monoamines and behavioral patterns in fish model system

**DOI:** 10.1101/2022.07.25.501403

**Authors:** Evgeny Ivashkin, Stefan Spulber, Andrei Zinovyev, Takashi Yoshitake, Shimako Yoshitake, Olga Kharchenko, Marina Yu. Khabarova, Spyridon Theofilopoulos, Jan Kehr, Ernest Arenas, Sandra Ceccatelli, Elena E. Voronezhskaya, Igor Adameyko

## Abstract

The presence of serotonergic system during early pre-neural development is enigmatic and conserved amongst all studied invertebrate and vertebrate animals. We took advantage of zebrafish model system to address what is the role of early serotonin before first neurons form. Unexpectedly, we experimentally revealed the existence of delayed developmental neurogenic and behavioral effects resulting from the manipulations of pre-neural (zygote, blastula and gastrula) serotonergic system. In particular, the delayed effects included differences in the synthesis of serotonin in early serotonergic neurons in the central nervous system as well as in behavioral alterations after habituation in zebrafish larvae. These effects appeared as highly specific and did not coincide with any major abnormalities. The same manipulations of the serotonergic system at neural developmental stages did not show such effects, which confirms that early effects of serotonergic system manipulation are not based on retained serotonin in embryonic cells. Accordingly, gene expression analysis demonstrated specific changes only in response to the elevation of early pre-neural serotonin, which included the delayed and pre-mature onsets of different gene expression programs. Taken together, our results introduce a novel function of early pre-neural serotonergic system in a vertebrate embryo – tuning and fine control of specific mechanisms at later neural developmental stages that result in a mild variation of a behavioral adaptive spectrum.

## Introduction

Small molecules represent one of the most ancient and abundant groups of regulators spread in all branches of life. Presumably, appearing as metabolites of structural and energy-carrying molecules, many of those underwent long evolution diversifying their roles in nature. Serotonin (5-hydroxytryptamine, 5-HT) is one of those ubiquitously-spread regulatory small molecules. Indeed, 5-HT is a well-recognized component of many ancient and archetypical signaling systems widely distributed among all major phyla [1].

In vertebrates, 5-HT serves as a classical neurotransmitter and a hormone playing a key role in diverse cellular and physiological processes such as memory formation, feeding, social and reproductive behavior, smooth muscle contraction, ciliary motility, blood coagulation and many others [2].

5-HT is synthesized from tryptophan and requires two enzymes that must act sequentially: the rate-limiting enzyme tryptophan hydroxylase (TPH) followed by aromatic L-amino acid decarboxylase (AADC). The teleost fish, a popular model system for investigating the serotonergic system, has two genes of TPH, two genes of 5-HT membrane transporter (SERT) and one gene of monoamine oxidase (MAO). MAO is responsible for 5-HT catabolism that also serves as a source of reactive oxygen species implicated in cell signaling [3]. There are exceptions to the rule and some teleost fish, for instance, zebrafish and medaka, have three forms of TPH (*tph1a, tph1b* and *tph2*). In addition, tyrosine hydroxylase (TH2) can also synthesize 5-HT in zebrafish [4]. Concentration of 5-HT inside of a cell may rely on a speed of SERT-mediated 5-HT re-uptake, which is K^+^/Na^+^-dependent and can be controlled by phosphorylation, NO-synthase and integrin coupling [5–7]. Importantly, some 5-HT-dependent regulations are based on serotonylation - a covalent binding of 5-HT molecule to a target protein by transglutaminase [8], which renders intracellular localization and local concentration of 5-HT an important parameters. Most of the 5-HT effects are implemented through a number of membrane-bound receptors that are subdivided into seven main families. Numerous genes of these 5-HT receptors are duplicated in teleost fishes, which led to diversification of their functions as compared to their homologs in other vertebrates [9].

Remarkably, even the organisms lacking nervous system such as bacteria, protists, plants, and some multicellular animals are also capable of producing and using 5-HT. In line with this, all animals with nervous system, where serotonergic system was investigated in the past, revealed the presence of 5-HT in oocytes, cleaving blastomeres and early pre-neural developmental stages, where the role of 5-HT is not directly related to the nervous system development and is currently poorly understood. When it comes to diverse and enigmatic functions of the early pre-neural serotonin, it appeared that 5-HT can stimulate or block maturation of oocytes as well as it can take part in the re-initiation of meiosis in a number of animals [10–12]. Mutations in 5-HT synthesis genes in *Drosophila* result in malformations of embryonic cuticle and can also lead to death of an embryo during pre-neural stages [13]. In addition, application of specific antagonists of 5-HT receptors leads to a complete block or, at least, a significant slowdown of cleavage divisions in nudibranch mollusk and sea urchin embryos [14,15]. In a freshwater gastropod snail *Lymnaea stagnalis*, the increase of 5-HT levels at early cleavage stage results in later disruption of gastrulation [16]. In amphibians and birds, 5-HT and its proper distribution within a cleaving embryo is an important condition for the left-right polarity-establishing mechanism [17–20]. Finally, early pre-neural embryos of the teleost fish show the clear presence of 5-HT, while the role of serotonin at such early stages was not sufficiently investigated [21,22]. In line with this, 5-HT receptors HTR1aa, 1ab, 1b, 2a and 5a, turned out to be expressed in a zebrafish cleaving and gastrulating embryos [23]. At later, neural developmental stages, 5-HT signaling is involved in pharyngeal arch morphogenesis in zebrafish [21], *Xenopus* [24] and mice [25]. In the nervous system of teleost fish and mammals, 5-HT promotes developmental [26,27] and adult neurogenesis [28]. Depletion of 5-HT in zebrafish larvae affects locomotor behavior as well as the body length and notochordal morphology [29]. Thus, all findings mentioned above suggest that 5-HT is an important integrative regulator of a fish development combining traits that are common with other vertebrates.

Notably, serotonin clearly participates in a variety of reproductive functions in the adult fish including the control of gonadotropin-releasing hormone, gonadotropin and luteinizing hormone release, gonadal maturation, and socio-sexual behavior [30]. More generally, 5-HT has been identified as an important regulator of fish female gonadal function [31–34] and a modulator of the effects of melatonin on steroid-induced oocyte maturation [35]. Furthermore, 5-HT is known as an important neuro-endocrine regulator of sexual plasticity [36], and the increased serotonergic signaling can prevent sex change in the saddleback wrasse in socially permissive conditions [37]. Such specific involvement in reproduction, maturation of gonads and production of oocytes together with the presence at early pre-neural embryonic stages suggests that the mother-derived serotonin can influence some aspects of embryonic development and, through this, can transmit non-genetic information from parents to progeny. Recently, we revealed such a case – a novel role of serotonin in a gastropod mollusk development, behavior and ecology, where 5-HT mediated non-genetic information transmission based on serotonin production in the reproductive tract of a mother snail [38]. This mother-derived 5-HT influenced the development of the snail embryo and caused delayed behavioral effects in juvenile individuals.

In this study, we tested the possible evolutionary-conserved information-transmitting role of pre-neural serotonin in a vertebrate teleost model system using zebrafish in a number of pharmacologic, embryologic and behavioral experiments. As a result, we demonstrated that early pre-neural 5-HT influences the later stages of neural development in a systemic way, and also leaves a footprint on the behavior of juvenile fish larvae, thus, making this mechanism of 5-HT-based adaptive information transfer possible in nature.

## Results

### 1. Pre-nervous serotonergic system in zebrafish is in place and active

To investigate pre-nervous serotonergic system, we firstly analyzed the presence and distribution of 5-HT by immunohistochemistry (Fig. 1A-H). Serotonin itself was expressed at all developmental stages analysed (from 2-cell to 50% epiboly). We found that 5-HT was present in the early zebrafish embryos mostly in a blastodisc and later in a blastoderm, being equally distributed both in a cytoplasm of each cell and between blastomers without any significant gradients (Fig. 1 A-C, F-H). Brightness of anti-5-HT staining, and thus expression, compared to the cytoplasm was higher in the nuclei of some cells (Fig. 1C). Most of the staining showed distinct punctate pattern corresponding to small cytoplasmic vesicle-like structures 300-700 nm in diameter in blastoderm, blastodisc and in periblast, but not in yolk (Fig. 1D, E). After the formation of a periblast, 5-HT was present there in the same concentration as compared to the nearby cells, while forming a sharp drop in expression at the border with yolk (Fig. 1D, G).

**Fig. 1.**
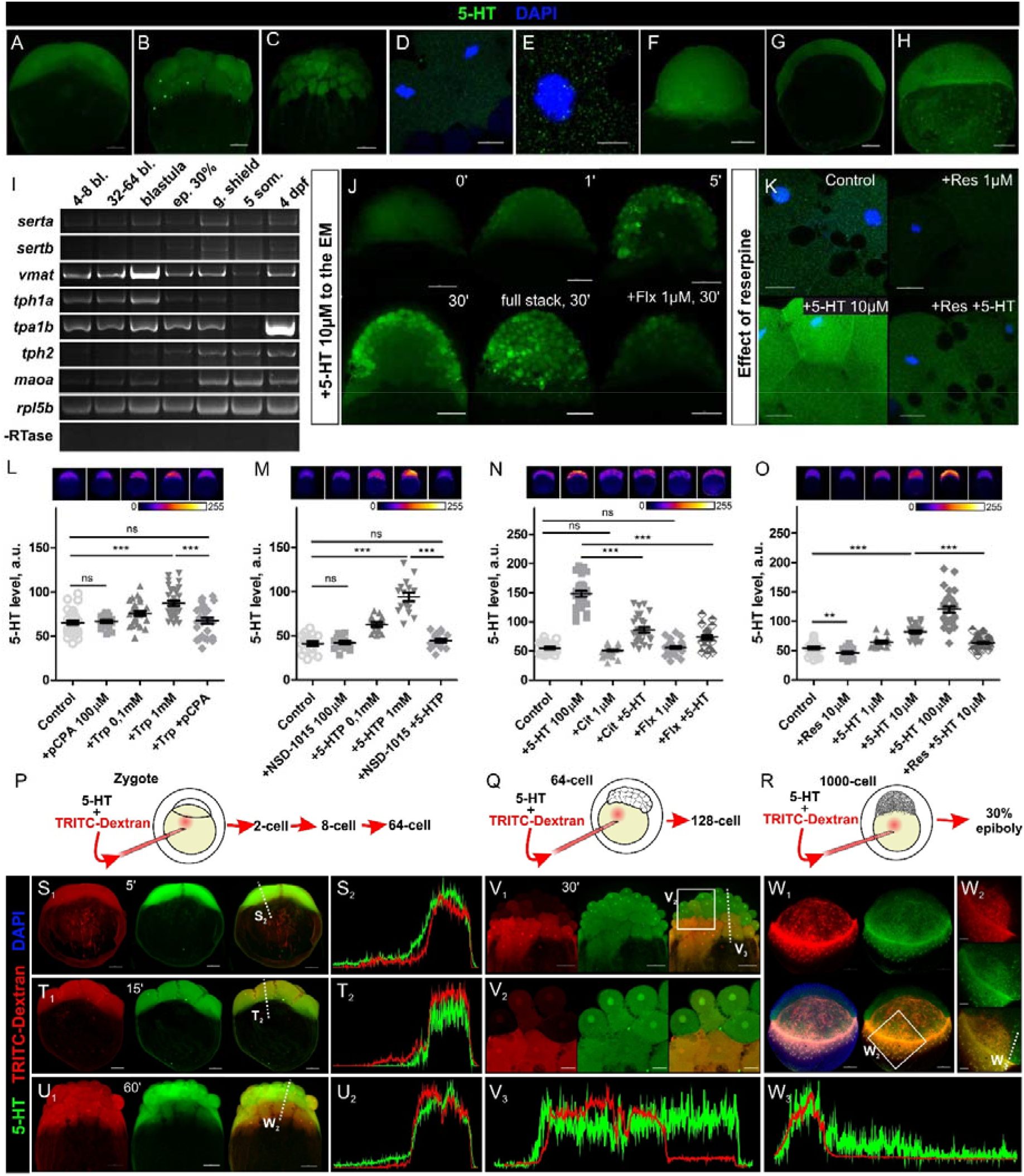
Distribution, synthesis and transport of 5-HT in zebrafish embryo during early developmental stages. (A-H) – Distribution of 5-HT in zebrafish early embryos. (A) – 2-cell stage; (B) – 32-cell stage; (C-E) – 128-cell stage; (F) – blastula; (G-H) – 50%-epiboly. (A-C, F and H) – General view. (D-E, G) – Optical sections. Note the absence of a 5-HT concentration gradient within the cleaving part of the embryo. (I) – Expression of 5-HT membrane transporters (serta, sertb), vesicular monoamine transporter (vmat), tryptophan hydroxylases (tph1a, tph1b, tph2) and monoamine oxidase (maoa) and ribosomal protein rpl5b as an internal control in embryos and 4dpf larva. (J) – Zebrafish embryo (blastula) is capable to uptake 5-HT from the media (for other stages see Fig. S1). Sagittal optic sections and a full stack (30 min) of control and experimental embryos from three consecutive time points after 5-HT application. Fluoxetine (Flx) selectively inhibits the 5-HT reuptake. Note the variation in 5-HT uptake capability among cells both in periderm and in deeper layers. (K) – Reserpine, a specific blocker of monoamines vesicular transporter, causes a decrease of 5-HT level. High magnification images, single optical sections of 128-cell embryo. (L-O) – Quantification of a 5-HT level in the cleaving part of the embryo after specific pharmacological manipulations. Relative brightness of a 5-HT immunostaining is plotted on the corresponding graphs. 1^st^ and 2^nd^ steps of 5-HT synthesis (tryptophan hydroxylase (L) and aromatic decarboxylase (M), respectively) as well 5-HT uptake (N) and release (O) are tested. Note that inhibition of neither 1^st^ nor 2^nd^ stage of 5-HT synthesis results in significant 5-HT decrease. But the same substances reduce the synthesis of 5-HT from supplemented precursors. SSRIs (fluoxetine and citalopram) demonstrate the same effect on 5-HT level within the analyzed cells. While blocker of vesicular monoamine transport reserpine reduces a 5-HT level. Data represented as mean ± s.e.m.; T-test; n=20-60 embryos; ns – non-significant; ** – p<0.01; *** – p<0.001. (P-W) – Analysis of 5-HT distribution after co-injection with TRITC-dextran marker. (P-R) – Experimental strategy outlines. (S1, T1, U1, V1 and W1-W2) – maximum intensity projection view; (V2) – optical section. (S2, T2, U2, V3, W3) – Linear brightness plots for the green and red channels generated from a single optical section. Note that 5-HT localization and amounts do not differ from the uniform distribution of 5-HT in a later embryo after the injection in a zygote or in a 64-cell embryo. At the same time, the injection into blastula leads to the preferential accumulation of 5-HT in the periblast with some gradient forming in the adjacent cells. Scale bars: (A-C, G-H, J, S1, T1, U1, V1, W1) – 100 μm; (D, K) – 15 μm; (E) – 5 μm; (V2) – 30 μm; (W3) – 50 μm.

Expression of different components of serotonergic system, including enzymes of 5-HT synthesis and degradation, 5-HT membrane and vesicle transporters was addressed by PCR at subsequent developmental stages of zebrafish. *Tph1b* (*tryptophan hydroxylase 1b*), *serta* (*slc6a4a, serotonin transporter SERTa*), *vmat* (*slc18a2, vesicular monoamine transporter*) and *maoa* (*monoamine oxidase a*) appeared to be expressed at pre-nervous stages of 4-8 blastomeres, 32-64 blastomeres, blastula, 30% epiboly, germ shield stage and 4dpf (Fig. 1I). Expression of *tph1a* (*tryptophan hydroxylase 1a*) was detected at aforementioned stages while expression of *tph2* (*tryptophan hydroxylase 2*) and *sertb* (*slc6a4b, serotonin transporter b*) was clearly observed only from blastula stage and onwards (Fig. 1I).

In addition to direct PCR we analyzed zebrafish and human developmental transcriptomics data obtained from independent studies [39,40]. We found an evident gene cluster containing transcripts related to 5-HT and its derivative melatonin that demonstrated the expression maxima specifically at pre-neural stages of zebrafish development. This cluster of genes included transcripts essential for the synthesis of these compounds, their catabolism, transport, receptors and some downstream genes (Fig. S1A). This notion was found to be conserved in other vertebrates including humans (Fig. S1B).

To investigate the activity of serotonin synthesis system we performed functional tests based on the incubation of zebrafish embryos at 16-32 blastomeres, 64-128 blastomeres, blastula and germ shield stages in biochemical precursors of 5-HT. Incubation of embryos at all aforementioned stages in L-tryptophan and 5-hydroxytriptophan resulted in a statistically significant increase in brightness of anti-5-HT staining (Fig. 1L, M; Fig. S2A, C-F). On the contrary, intensity of staining in yolk did not show significant changes in response to pre-incubation in those compounds at all analyzed stages (Fig. S2B). Interestingly, pharmacological blockade of both key synthesis enzymes did not result in a drop of the 5-HT level. At the same time, incubation of early embryos in 5-HT biochemical precursors with the simultaneous 5-HT synthesis blockade, resulted in lower 5-HT levels compared to when embryos were incubated only in biochemical precursors (Fig. 1N, O).

Both uptake and efflux of 5-HT are active processes that play an important role in maintaining proper balance of 5-HT concentration in the developing embryo. To address the uptake of serotonin at pre-nervous stages we firstly incubated teleost embryos in solutions with a range of 5-HT concentrations. We identified that all cells in cleaving and gastrulating embryos, could efficiently uptake 5-HT in a concentration-dependent mode (Fig. 1N, O; Fig. S2A-G). In case of blastula, after the 5-HT uptake, the gradient of 5-HT formed between the outer and the inner cell “layers” of the blastoderm (Fig. 1J). As expected, SSRIs could decrease the 5-HT uptake, whereas none of these compounds significantly influenced natural 5-HT levels (Fig. 1N). Treatment with reserpine, a VMAT inhibitor, decreased 5-HT levels via allowing cytoplasmic vesicles to release 5-HT (where it is catabolized by MAO) while blocking the re-loading of the vesicles with 5-HT via VMAT action (Fig. 1K, O). The effect of injection of reserpine was similar to the effect of incubation of embryos with reserpine solution. Reserpine treatment also decreased 5-HT level in case of a co-incubation of embryos with 5-HT (Fig. 1K).

In order to assay intracellular transport of 5-HT and the effects of disturbing 5-HT distribution in embryo, we injected 5-HT into yolk and blastomeres. After the injections of 5-HT into the yolk from zygote to 32-cell stage, 5-HT efficiently moved into the cytoplasm of blastomeres. Administered 5-HT left only slight traces in the yolk already after 5 min post-injection (Fig. 1P, S_1_-S_2_) and absolutely no traces were observed in 15 min and later (Fig. 1T_1_-U_2_). At the same time, the co-injected TRITC-dextran (10 KDa) was also distributed evenly from zygote to 16-cell-stage but formed a lasting gradient in between cells in case of being applied at 32-64-cell stages (Fig. 1Q, V_1_-V_3_). Also no 5-HT gradient was formed in case of 5-HT injections into separate blastomeres at least up to 32 cell-stage. 5-HT staining was equalized between blastomeres already in 10 min after injection (data not shown). At a later timepoint, after the injection into a yolk cell at a blastula stage, most 5-HT stayed in the periblast. A weak gradient of 5-HT formed in nearby cells, but it was quickly dissolved in a layer of 10-15 cells (Fig. 1R, W_1_-W_3_). When the injection of 5-HT was made into a zygote, increased anti-5-HT staining brightness was retained in an embryo and could be detected with immunohistochemistry at least up to 32 hpf (Fig. S2I).

### 2. Early pre-nervous serotonergic system modulation causes delayed effects in larvae

We firstly investigated larvae at 1, 2 and 4 dpf after anti-5-HT immunostaining. At 1 dpf larvae injected with 5-HT into zygote demonstrated increased levels of 5-HT immunoreactivity in all tissues and even in the lumen of the neural tube. However, we could not identify 5-HT positive neurons in the brain and neural tube from these injected animals as compared to the non-injected controls (Fig. 2A_1_-A_4_, C_1_-C_4_). In line with this, in these experimental animals, anti-5-HT staining in the serotonergic sensory cells in the skin was poorly contrasted. When we checked the 2 dpf larvae injected at zygote, we consistently observed the reduced contrast of 5-HT staining in serotonergic neurons of hypothalamus, anterior and inferior raphe, arrowhead population and pineal gland (Fig. 2H_1_-I, O-T). At 4 dpf, 5-HT immunostaining in the CNS of treated embryos did not differ from controls. Importantly, the 1-2 dpf larvae injected with 5-HT at blastula stage (contrary to the injection at zygote stage) did not show any differences to control embryos in staining of serotonergic cells while could demonstrate some elevation of 5-HT level in some tissues (mostly in trunk; Fig.2D_1_-G).

**Fig. 2.**
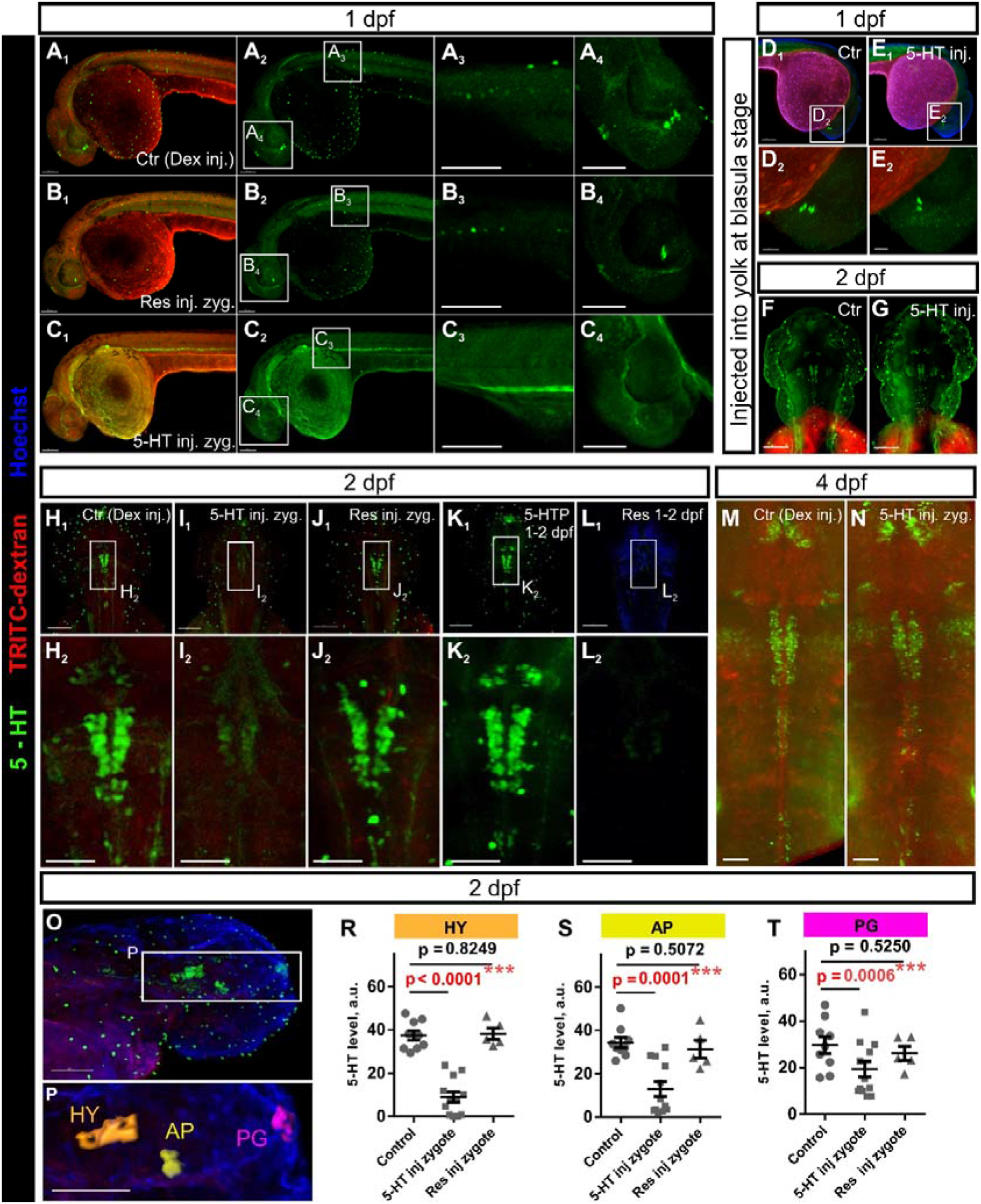
Modulation of pre-neural 5-HT causes delayed effects in 5-HT distribution in zebrafish larvae. (A-C) – 5-HT-positive cells in 1 dpf zebrafish larva injected with 5-HT or reserpine in combination with TRITC-dextran at zygote stage. (A1-A4) – TRITC-dextran- and DMSO-injected control; (B1-B4) – injection with reserpine; (C1-C4) – injection with 5-HT. Note that in case of 5-HT administration, the levels of the background staining in tissues are increased, while characteristic 5-HT-containing cells are not contrasted at all. (D-G) – 5-HT-expressing cells in the head region of the fish injected with 5-HT and TRITC-dextran into yolk at a blastula stage. (D-E) – 1 dpf, (F-G) – 2 dpf. (D1-D2, F) – TRITC-dextran and DMSO-injected control; (E1-E2, G) – 5-HT-injected fish. (H-L) – 5-HT-expressing cells in the head of 2 dpf zebrafish larvae. (H1-H2) Vehicle; (I1-I2) fish injected with 5-HT and (J1-J2) reserpine at a zygote stage; (K1-K2) fish incubated in 1 mM 5-HTP and (L1-L2) 10 μM reserpine for 24 hours between 1 dpf and 2 dpf. (M-N) – 5-HT-positive neurons in the brain of 4 dpf larvae. Vehicle (M) and fish injected with 5-HT at a zygote stage. (O-T) – Relative brightness of the anti-5-HT staining in cells of 2 dpf larvae. (O, P) – head of the fish (O) with 3D rendering (P) used to generate measurements of brightness in the volume of serotonergic cells plotted on the graphs in panels (R-T). (HY) - Cells belonging to hypothalamus together with superior raphe; (AP) arrowhead population and (PG) pineal gland. (R-T) – brightness of 5-HT staining measurements. Data represented as mean ± s.e.m.; T-test; n=7-10 larvae. Scale bars: (A-C, D1, E1, F-G, H1, I1, J1, K1, L1, O-P) – 100 μm; (D2, E2) – 30 μm; (H2, I2, J2, K2, L2) – 50 μm.

In order to obtain positive control, we incubated embryos in 5-HT precursor 5-HTP at early neuronal developmental stages (between 1 and 2 dpf). Later (2 dpf and beyond) these larvae demonstrated generally elevated levels of 5-HT in numerous tissues as well as in serotonergic neurons located in CNS. While all serotonergic elements persisted well distinguishable in staining (Fig. 2K_1_-K_2_).

Contrary of the experiments with 5-HT level exogenous elevation, both incubation and injections of reserpine into a zygote did not influence the level and localization of 5-HT at 1-4 dpf (Fig. 2B_1_-B_4_,J_1_-J_2_, O-T). On the contrary, incubation in reserpine at 1-2 dpf drastically decreased the level of 5-HT in all tissues (Fig. 2L_1_-L_2_).

Despite all direct manipulations with 5-HT levels in cleaving embryos, the general morphology of larvae was not affected notably (Fig. S3A). Since 5-HT depletion is known to affect growth of larvae [29] we assayed rostro-caudal length and head-to-body length proportion in larvae at 2 and 4 dpf after incubation in 5-HT and other substances from zygote to epiboly stage (Fig. S3B-F). We did not detect changes in length of the experimental fish after 5-HT (Fig. S3A) although some drifts of body proportions were noticeable. In case of reserpine incubations, experimental fish turned out to be shorter without body proportion change. The strongest effects on the fish morphology were observed after the incubations in the blocker of the 2^nd^ stage of 5-HT synthesis NSD-1015 and tryptophan. NSD-1015 and pCPA influenced the proportions of the head and the rest of the body, while tryptophan made fish proportionally shorter (scaling down the general size). In all these cases, the effects were rather minor but significant in case of 3-4 repetitions and large numbers of fishes analyzed (Fig. S3C-E).

In addition to the general morphology and growth, we analyzed possible left-right asymmetry anomalies by looking at cardiac morphology after incubation in blockers of 5-HT synthesis, reserpine, number of SSRIs and antagonist of 5-HT_4_ receptors GR-113808, which is known to affect normal left-right asymmetry in Xenopus embryos being applied at early developmental stages [18]. We did not observe any significant differences between experimental and control groups (Fig. S4B).

Furthermore, we did not find any visible disturbances of early development in the fish incubated between zygote- and 50% epiboly-stage in a number of known serotonergic system modulators we tested. Namely, antagonists of 5-HT receptors (1-100 μM): methiothepin, mianserin, metoclopramide, NAN-190, trazodone, ritanserin, SB 203186, SB 269970, spiperone, (S)-WAY100135, tropanil, zakopride; 5-HT receptor agonists (1-100 μM): 5-CT, 5-MeO-DMT, 8-OH-DPAT, alpha-methylserotonin, Br-LSD, DOI, mCPP; 5-HT transporter blockers (1-100 μM): fluoxetine, citalopram, fluvoxamine, imipramine, clomipramine. Blockers of the transglutaminase-mediated serotonylation MDC (100 μM) and cystamine (1 mM) also did not affect early development of fish.

### 3. Modulation of serotonin level at the early stages affects content of monoamines and their metabolites in larvae

In order to perform direct measurements of monoamines and their metabolites, we utilized a highly sensitive H PLC-based approach that allowed us to perform investigations at the level of individual fish embryos or larvae (or their separated heads) (Fig. 3A, S5A). The whole-body analysis showed that 2 dpf larvae injected with 5-HT at zygote stage demonstrate significant general elevation of 5-HT and its major metabolite 5-HIAA (Fig. 3B, E, H-I). At the same time, the proportion of 5-HIAA and 5-HT did not differ from control in the case of whole-body measurements. However, if heads were analyzed separately from the body, we detected differences in the proportion of 5-HT and its metabolite: the level of 5-HT did not differ significantly from control larval heads, while 5-HIAA levels turned out to be 1.5-fold lower than a norm. However, at 6 dpf the experimental fish heads showed similar 5-HT and 5-HIAA levels as compared to controls (Fig. 3D, F). Incubation of 1-2 dpf larvae in 5-HTP significantly increased the levels of 5-HT in the heads of 2 dpf larvae, while 5-HIAA was slightly (non-significant) increased (Fig. 3C, F). These experimental fish maintained the differences in 5-HT (non-significant elevation) and 5-HIAA (significant elevation) even at 6 dpf (Fig. 3D, F). Injections of reserpine into a zygote caused a small but significant elevation of 5-HT in the head at 2dpf larvae. Incubation of 1-2 dpf larvae in reserpine led to a cardinal reduction of all monoamines and also shifted metabolite/monoamine ratio towards metabolites (Fig. 3C-D, G).

**Fig. 3.**
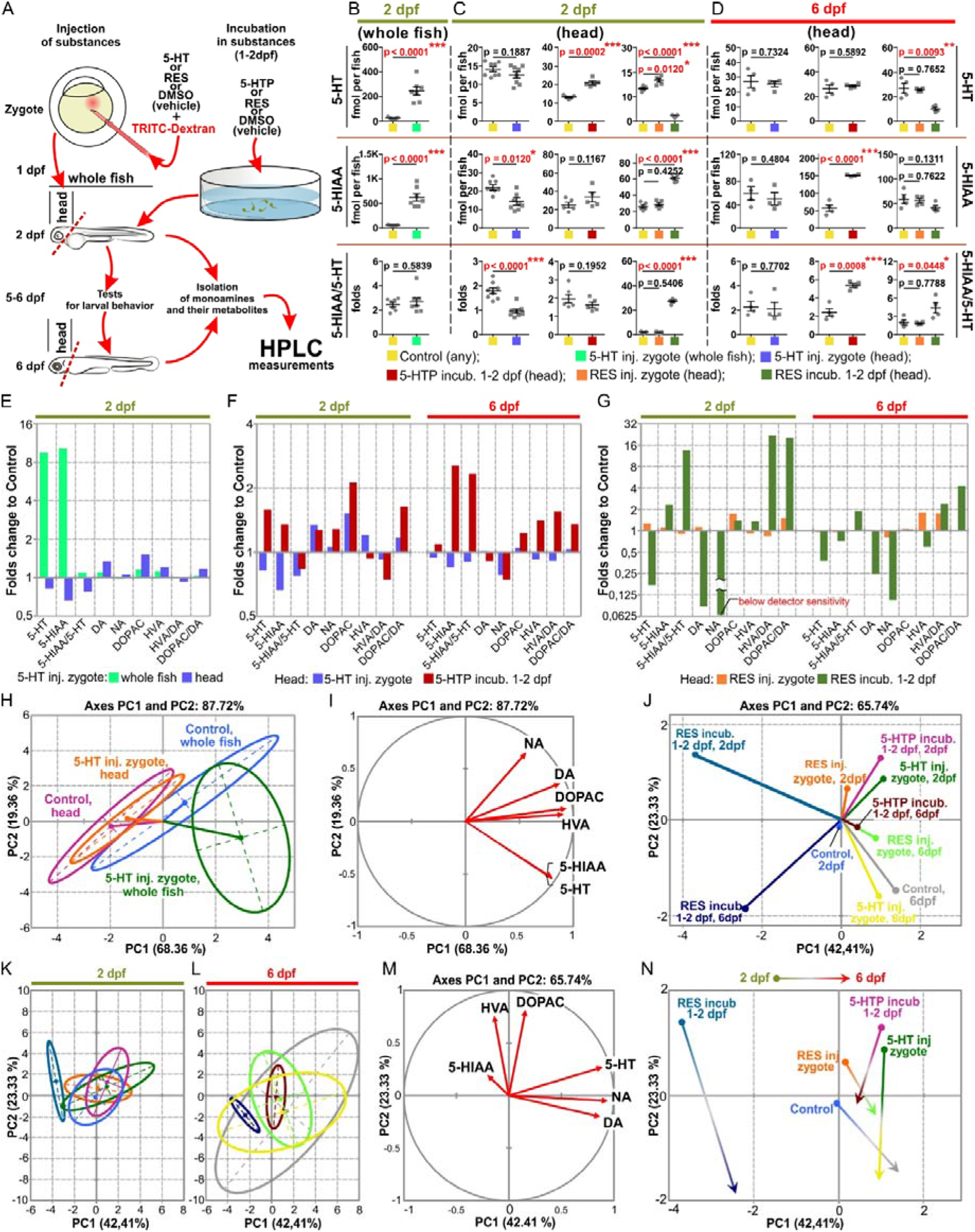
HPLC measurements of monoamines and their metabolites in fish after the modulation of 5-HT levels at pre-neural (zygote) and early neural (1-2 dpf) developmental stages. (A) – Scheme of the experiment. (B-D) – 5-HT and its main metabolite 5-hydroxyindoleacetic acid (5-HIAA) content in the whole 2 dpf zebrafish larvae (B) and in the separated heads of 2 dpf (C) and 6 dpf (D) fish. 5-HIAA/5-HT-metabolite to monoamine ratio. Measurements are plotted for the control (vehicle), the fish injected with 5-HT or reserpine at a zygote stage and for the larvae incubated in 1 mM 5-HTP or 10 μM reserpine for 24 hours between 1 and 2 dpf (see the color code in the figure). In (B-C) the measurements were done for 5 whole fish or severed heads per sample (B, C) and for the one fish in (D). Data for the other measured substances can be found in Fig. S5. Data represented as mean ± s.e.m., T-test. (E-G) – Summarized results of measurements of relative quantities of all analyzed substances shown as folds to control. Groups of experiments were the same as outlined in (B-D). (H-N) – PCA (principal component analysis) plots of monoamines and their metabolites content as variables identified in the experimental groups listed above. (H-I) – Comparing the monoamine levels in 2 dpf whole fish and separated heads in control and after 5-HT administration in a zygote. (H) – Group barycenters shown on the plane of the first two principal components together with 95% confidence ellipses. (I) – Visualizing variable contributions to the first two principal components of (H). (J-N) – Comparisons of the monoamine levels in 2 dpf and 5 dpf fish heads analyzed at different experimental steps. (J) – All group barycenters shown on the plane of the first two principal components. Confidence ellipses are plotted for the experimental groups at 2 dpf (K) and 6 dpf (L) derived from the same PCA plot. (M) – Visualizing variable contributions to the first two principal components of (J). (N) – Barycenter displacement for the 2 dpf and the 6 dpf groups shown on the plane of the first two principal components.

Since behavioral effects of 5-HT metabolism changes in brain are often promoted through the coordinated reflection in other monoamine systems, catecholamines noradrenaline (NA), dopamine (DA) and its metabolites DOPAC and HVA were also taken into analysis (Fig. S5B). DA and its main metabolite DOPAC level was increased in the heads of 2 dpf fish subjected to the elevation of 5-HT both at zygote and 1-2 dpf (through the application of 5-HTP) stages. By 6 dpf these differences disappeared. NA was affected only by later but not early increase of 5-HT. It was higher than in control in 2 dpf and lower in 6 dpf. DOPAC unlike HVA was touched in both variants of experiment with 5-HT increase.

Taken together, changes in monoamines and their metabolites differ in the head and whole body of fish. Principal components analysis reveals that changes in measured substances level after the injection of 5-HT into a zygote are different (and often opposite) from the changes in experimental fish incubated with 5-HTP at neural 1-2 dpf stages. Accordingly, incubation in reserpine at 1-2 dpf caused different effects on the concentration of serotonin and catecholamines as compared to the effects resulting from the injection of reserpine into a zygote. Generally, the effects of 5-HT injected at a zygote stage on later monoamine concentrations were almost undetectable at 6 dpf. On the other hand, the effects of incubation in 5-HTP at 1-2 dpf remained pronounced at 6 dpf (Fig. 3J-N).

### 4. Larval behavior is affected by increase rather than decrease of 5-HT in the early pre-nervous embryos

Disturbances in the level of 5-HT or catecholamines have been shown to cause specific changes in larval behavior of zebrafish [41]. In order to confirm our observation on early 5-HT action we performed number of behavior tests on 1 dpf – 6 dpf old fish (see scheme at Fig. 4A).

**Fig. 4.**
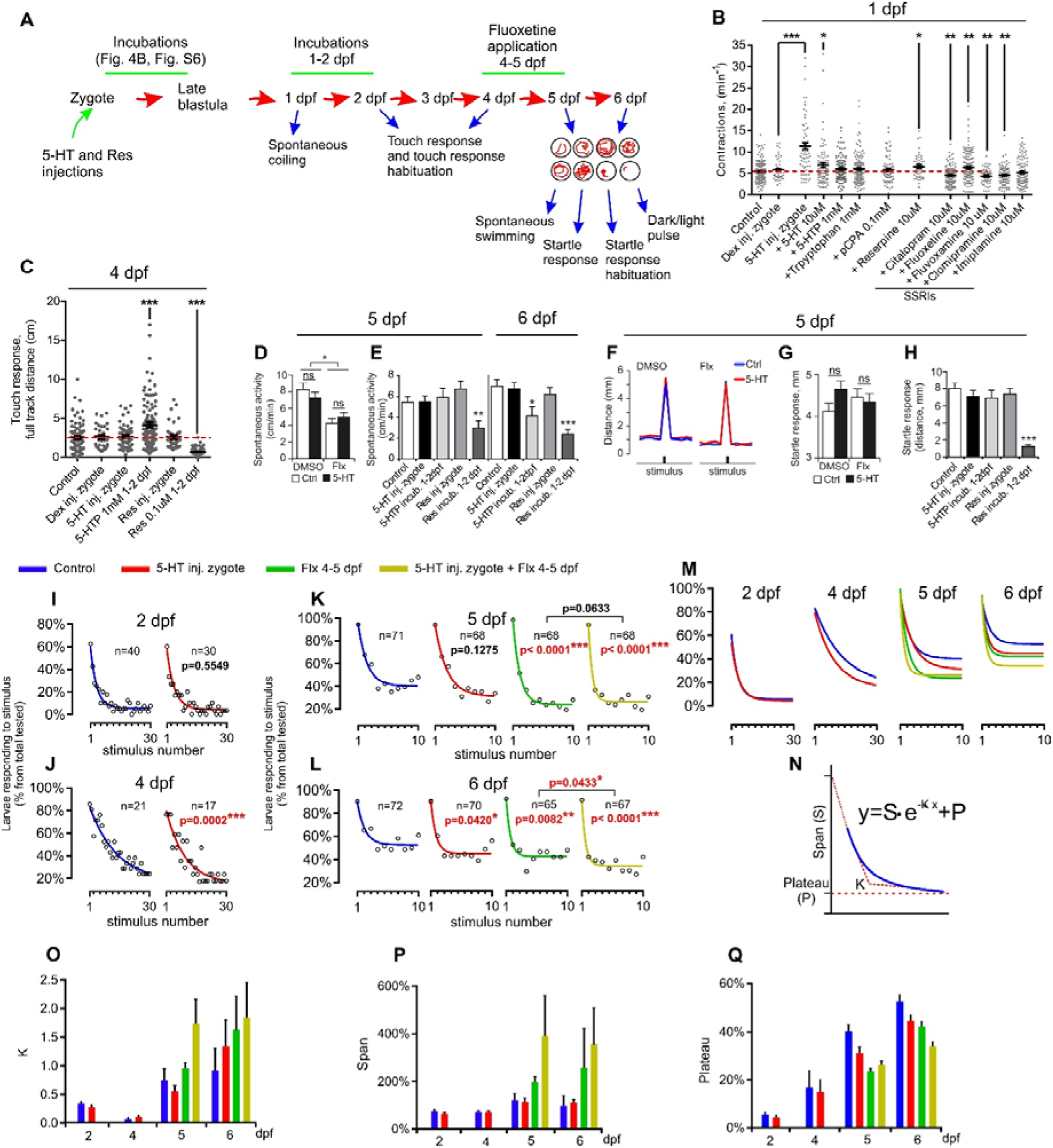
Selective modulation of a 5-HT level in a cleaving pre-neural embryo alters larval behavior. (A) – Outline of treatments and behavioral assays. (B) – The frequency of spontaneous coiling of a trunk in 1 dpf larvae after the 5-HT injections into a zygote and also after the incubation of embryos in different substances starting from a zygote till late blastula stage. (C) – Startle response to tactile stimulation, 4 dpf. (D, E) – Spontaneous swimming activity of 5 dpf and 6 dpf larvae. (D) – Larvae administered with 5-HT at a zygote stage without or with the following incubation in fluoxetine for 24 hours between 4 dpf and 5 dpf. (E) – Fish injected with 5-HT or reserpine at a zygote stage and either incubated in 1 mM 5-HTP or in 10 μM reserpine for 24 hours between 1 and 2 dpf. (F-H) – Startle response test (sound stimulation). (F) - Illustration of a startle response. (G) – The same design as shown in panel (D). (H) – The same design as shown in (E). (I-R) – Short term habituation to a touch response (I and J) and to a startle response (K and L). Results are presented as a proportion of the responding fish to the total number for each stimulus. (I and J) Graphs showing the data for the control (vehicle) condition and for the fish injected with 5-HT into a zygote. Plots in panels (K and L) represent the same treatments as described in a panel (D). Graphs show a non-linear regression with one phase exponential decay equation and p-values for the comparison of the regression equation parameters with an extra sum-of-square F-test. (M) – Summary plot for the regression graphs of the experimental points from panels (I-L). (N) – Explanatory graph for the equation parameters. X-axis is a time; Y-axis is a response. Y starts as equal to (Span + Plateau) and decreases to Plateau with a rate constant K. (O-R) – Comparison of fit’s equation parameters for the experimental points given on the plots (I-L). Span and Plateau are expressed in the same units as the Y-axis. K is expressed in the inverse manner of the units used for the X-axis (stimulus number). Data represented as mean ± s.e.m. (B-H), or mean ± 95% confidence interval (O-Q); (B, C, E, G, H) - T-test; (D) – one-way ANOVA and T-test; (I-L) – extra sum-of-squares F test for the equality of regression fit curves; (D-H) - n=150-200 larvae; ns – non-significant; * – p<0.05; ** – p<0.01; *** – p<0.001.

We firstly found that elevation of 5-HT at early pre-nervous stages led to an increase in the frequency of coiling (spontaneous trunk contractions) in 1 dpf embryos. Pharmacological compounds enhancing synaptic 5-HT signaling (reserpine and SSRIs) also influenced this parameter (Fig. 4B). Startle response to touch test at 4 dpf revealed no significant differences to control in fish that were injected with 5-HT during a zygote stage. Similarly, we found no effects in either spontaneous swimming or startle response to acoustic stimulation at 5-6 dpf (Fig. 4F-H, S6A-B).

In contrast, larvae incubated in 5-HTP at 1-2 dpf demonstrated higher activity in a touch response test at 4 dpf (Fig. 4C) and reduced spontaneous activity at 6 dpf (Fig. 4E). At the same time, in these fish we did not find any differences compared to the control in the frequency of coiling (1 dpf), spontaneous swimming and acoustic startle response at 5 dpf (Fig. 4B, C-H). Thus, the effects of 5-HT level increase through the application of 5-HTP between 1 and 2 dpf differ from the effects resulting from 5-HT increase at pre-neural stages.

Decreasing 5-HT level at cleavage stages by reserpine injection into the zygote did not induce significant effects in any tests we have assayed (Fig. 4C-H). In contrast, reserpine application at 1-2 dpf significantly reduced all types of larval behavioral activities (Fig. 4C-H).

We next assessed the habituation to repetitive tactile (2-4 dpf) or acoustic (5-6 dpf) stimulation by measuring the proportion of larvae responding to sequential stimulus application. Larvae injected with 5-HT at a zygote stage demonstrated significant differences from control at 4-6 dpf in both aforementioned tests (Fig. 4I-M). We observed significant differences in the speed of decline in response to repetitive stimulations (Span and K – Fig. 4N, O, P), as well as in the minimal proportion of responding fish (Plateau, Fig. 4N, Q). The differences in response to stimulation appeared to be larger at 6 dpf than at 2 dpf.

Serotonergic anterior raphe neurons in zebrafish are known to be implicated in the arousal and behavioral reactions to dark-light cycling [42] we decided to expose larvae to dark-light pulse test. The larvae were subjected to 10 min of complete darkness. Swimming activity was recorded continuously starting 10 min before the dark pulse, and for 10 min after the end of the dark pulse (Fig. S6C). The response to sudden dark onset is a complex behavioral reaction that involves multiple signaling systems, which eventually result in an abrupt increase in activity (about 1s), followed by sustained hyperactivity (Fig. S6C-D). Although the shape of the response appears similar (Fig. S6C, H, J), the larvae injected with 5-HT display a small but significant reduction in the initial response (Fig. S6I-J), hyperactivity decline in the darkness (Fig. S6C, H) and increased activity after light is switched on again (Fig. S6C, E, J-K).

In summary, our results revealed that the effect of a zygotic injection of 5-HT at zygote stage and incubation with fluoxetine at 4-5 dpf had similar effects. But expression level of the treatment’s behavioral effect in case of 5-HT administration to zygote was much milder than hyperserotonic phenotype resulted from SSRI treatment at 4-5 dpf. Incubation in fluoxetine at 4-5dpf significantly enhanced the effects of zygotic 5-HT injection (Fig. 4I-L; Fig. S6G, H, G-H). However, some parameters and reactions did not sum up (Fig. 4D, F-G; Fig. S6A-C, J). That indicates complexity of the effect and some differences to classical serotonin syndrome caused by SSRIs.

### 5. Exogenous elevation but not reduction of 5-HT in cleaving embryos induces systemic changes in the transcriptome of the developing zebrafish larva

Transcriptomic analysis in response to 5-HT increase and decrease at zygote stage was performed at 2 and 4 dpf in two replicates with control. The overall effect of 5-HT moderation on transcriptome was relatively modest compared to the control experiment: 2.5% of all probesets changed their expression more than two-fold in 2 dpf, and 1.3% of all probesets had more than two-fold change at 4 dpf, and only 0.2% of all probesets changed their expression more than 4-fold at 2 or 4 dpf compared to the control (Fig. 5B). Effect of 5-HT decrease by early reserpine injection was very moderate unlike to increase (Fig. 5C).

**Fig. 5.**
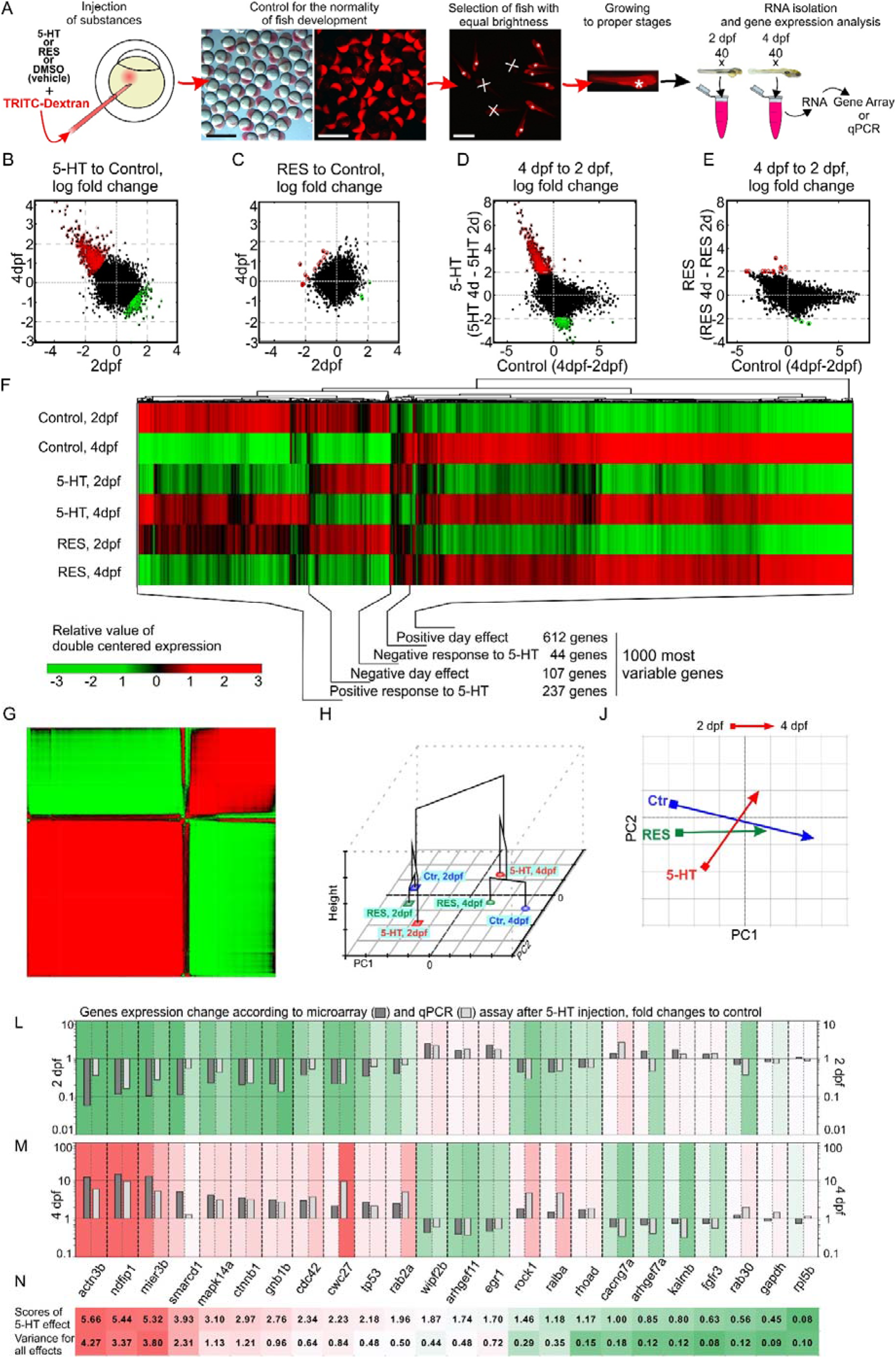
Increase of pre-neural 5-HT induces systemic and long-lasting changes in the transcriptome of the developing fish. (A) – Scheme of the experiment. (B, C) – Dependence of normalized response in 2 dpf and 4 dpf fish to 5-HT (B) and reserpine (C) injections at the zygote stage. For 5-HT Pearson correlation coefficient r = −0.41, p-value < 10^−16^. Red circles (for 5-HT n=611; 0.4% of the total number of probe sets) represent the probe sets with the log_2_ ratio between days 4 and 2 more than 2.0, and green circles (for 5-HT n=305; 0.2%) are the probe sets with the log_2_ ratio between days 4 and 2 less than −2.0. (D, E) – Day-effect on the response to 5-HT (D) and reserpine (E) as a function of the day-effect in the control experiments. (F) – Results of hierarchical clustering using 1000 most variable probesets across all conditions. (G) – Correlation matrix for the same probesets. For (G and H) the data were normalized by applying double centering to the initial logged expression matrix. (H) – PCA plot (horizontal plane) and clustering the experimental conditions (Z axis), based on 1000 most variable probesets; biological replicas are averaged. (I) – same PCA as on (H), shifts between day points are shown. (L, M) – Measured expression changes for the selected genes in 2 dpf (L) and 4 dpf (M) fish injected with 5-HT. The fold changes measured by microarray (dark grey) are validated by qPCR measurements (light grey). (N) – Scores of the 5-HT effect and variance among all experimental groups for genes chosen for qPCR validation.

In order to functionally characterize the effect of the serotonin and distinguish it from the transcriptome changes caused by development course itself, we considered three log ratios for the analysis: relative to control effect of 5-HT at 2 dpf (HT2), relative to control effect of 5-HT at 4 dpf (HT4), relative difference of gene expression in control experiments between 4 and 2 dpf (DAY_CONTROL). We observed striking negative dependence between normalized responses to serotonin in 4 and 2 dpf (Fig 5B). The difference in response to 5-HT elevation between 4 and 2 dpf was specific and not redundant with changes between days in control experiments (Fig 5D). We have not observed this effect for reserpine experiment, where the amplitude of the response was much smaller and did not show significant correlation between 2 and 4 dpf (Fig 5C, E). See the data for all probesets and recognized genes in the tab All probesets in Supplementary File 1.

Clustering analysis of 1000 most variable genes across all conditions revealed 4 major groups of genes (Fig. 5F, G): positive or negative 5-HT effect and positive or negative day effect. Importantly, the response to reserpine was substantially weaker and at the same time opposite to the response to 5-HT for the majority of genes in these identified groups where the input into the dispersion of response to the compound was higher as compared to the input into the dispersion of the day effect (Fig. 5F). According to the results of clustering analysis and PCA, the embryos injected with reserpine took an intermediate position between controls and embryos injected with 5-HT (Fig. 5H-I).

In order to validate microarray transcriptomics data on bulk RNA probes isolated from control and 5-HT-injected embryos, we selected a number of genes with different 5-HT scores and all-effects variance. qPCR assay confirmed that for the majority of these genes, the direction of their change and the order of their response correspond to the microarray data (Fig. 5J-L).

After ranking the genes according to their relative expression between the two days (HT4-HT2), we checked for the functional enrichment of zebrafish Gene Ontologies in the resulted ranking, using Gene Set Enrichment Analysis (GSEA). In the negative part of the ranking, we observed significant enrichment (corrected p-value<0.05) of several functional categories of genes related to the small GTPases: such as GO:0005083, SMALL GTPASE REGULATOR ACTIVITY (with wasb, cdc42bpb, mink1 in the top ranked genes in the core enrichment), GO:0051056, REGULATION OF SMALL GTPASE MEDIATED SIGNAL TRANSDUCTION (sos2, rgl3a, tsc2, rgl2 genes) and GO:0005452, INORGANIC ANION EXCHANGER ACTIVITY (with slc4 gene family in the top contributing genes) (See Supplementary File 1). These functional categories are upregulated compared to the control in day 2, and become relatively downregulated in day 4. The positive part of the ranking shows the opposite trend (downregulation in 2 dpf and upregulation in 4 dpf). In this part, we observed enrichment in diverse functional categories) such as GO:0006457, PROTEIN FOLDING (cwc27, sec63, vbp1, ppiab, uri1, dnajb11, fkbp9 in the top genes), GO:0016874, LIGASE ACTIVITY (ube2g1b, adssl, traf6, ube2na, arih1l, arih1, ube2a) and GO:0006184, GTP CATABOLIC PROCESS (rab2a, ralba, rhoad, nras). The detailed results of GSEA analysis are provided in the tab GSEA in Supplementary File 1.

In addition, we used ZEOGS tool for GO term analysis with a focus on anatomical categorization [43] which showed that for 2-4 dpf CNS was the most prominent GO term (tegmentum, CNS, cranial ganglion, hidbrain, forebrain, spinal cord and other). Also, the effect of 5-HT was apparent among the genes associated with peripheral sensory structures, cardiovascular and excretory systems. These effects were the most stable among nervous and excretory systems (Fig. 6A-E). See the full list of anatomical GO terms in the tab Anatomical GO Terms in Supplementary File 1.

**Fig. 6.**
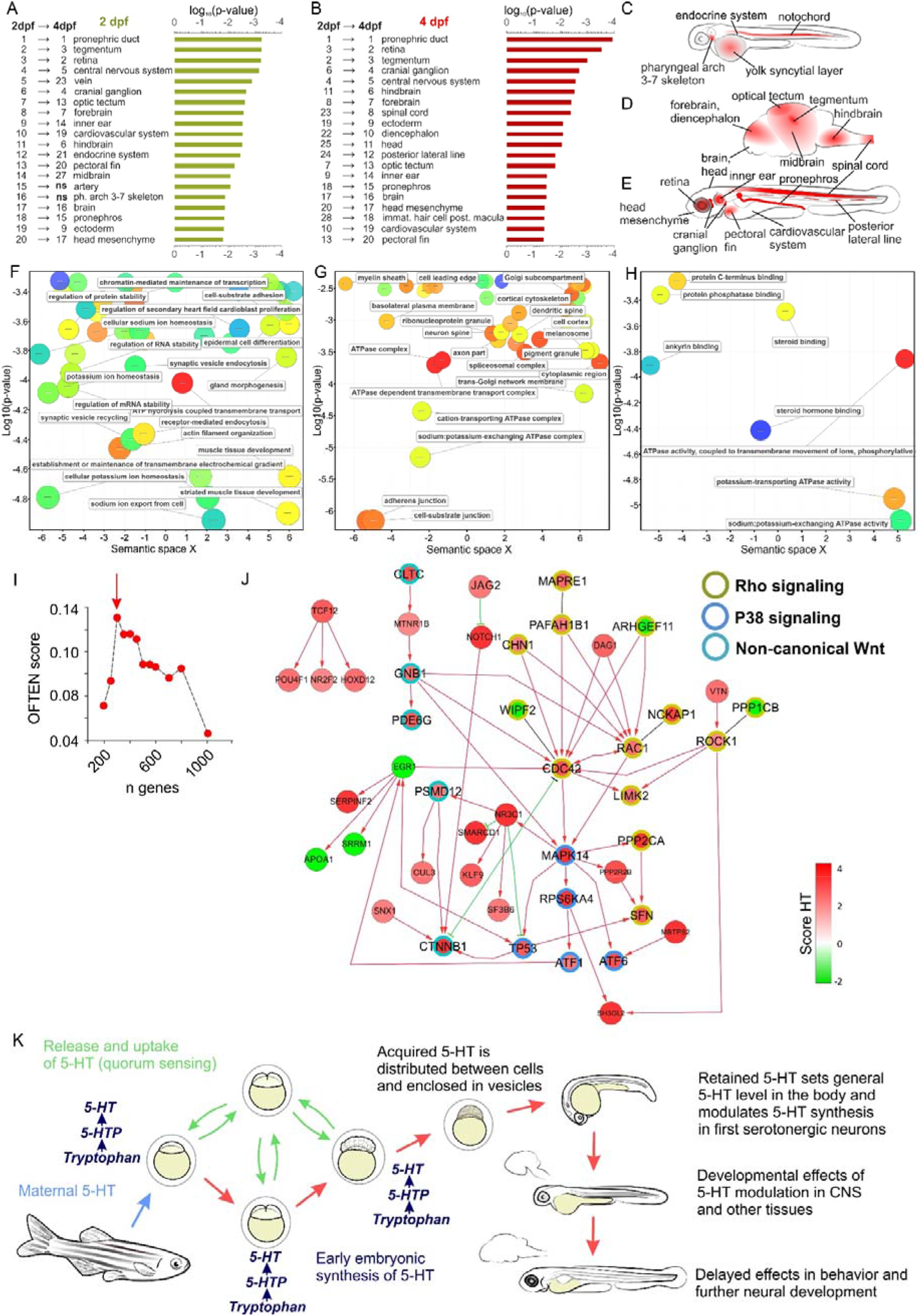
Analysis of systemic changes in the transcriptomes of zebrafish larvae in response to the early pre-neural 5-HT modulation. (A-E) – Gene ontology term enrichment analysis using ZEOGS tool for identifying organs characterized by specific expression of genes with the highest 5-HT-effect scores in 2 dpf (A) and 4 dpf (B) larvae. (C-E) highlights the locations corresponding to the most statistically significant anatomical terms for the genes in 2 dpf fish (C), 4 dpf fish brain (D) and whole 4 dpf fish (E). (F-H) – REVIGO visualization of gene set enrichment analysis (GSEA) results using gene ranking accordingly to the 5-HT-effect score. GO terms are grouped by biological process (F), cellular structure (G) and molecular function (H) of related gene sets. See Supplementary Dataset 1 for the complete lists. (J-K) – Network analysis of the genes with the highest level of 5-HT-effect scores, using network of high-confidence (not only computationally predicted) functional interactions from REACTOME pathway database (Zebrafish subset). (J) - top 5-HT-reacting genes are functionally related (interact with each other), more frequently than expected by random choice of genes with the same connectivity distribution; the significance is estimated using OFTEN score, with a peak at 300 top ranked genes. (K) – the functional interaction network between top ranked 5-HT-reacting genes, constructed from REACTOME; three over-represented pathways are highlighted by node edge color. (L) – Proposed model for the early 5-HT role in zebrafish.

Meta-analysis of GO term lists categorized according to the biological processes revealed membrane transport and ion homeostasis as the most robust groups. GO terms focused on the cellular structure showed the potential importance of a cell membrane as well as adhesive and cell-substrate junctions (Fig. 6F-H).

Next, we analyzed the potential reactome. Using OFTEN analysis tool, we found that the genes from the 5-HT score list formed a statistically significant network (Fig. 6I) with robust guanine-binding, RHOC-related and b-catenin-related subnets (Fig. S7). Among main hubs of reactome network we found such an important for the process of development genes as cdc42, rac1, egr1 mapk14 and ctnnb1. Application of analysis based on REACTOME pathway database [44] (zebrafish subset) revealed the highest score amongst the predicted networks connected with Rho, p38 and non-canonical Wnt signaling (Fig. 6J).

Next, we applied network-based analysis of top HT4-HT2 genes using REACTOME pathway database [44] (zebrafish subset). Using OFTEN analysis tool [45], we found that the first 300 genes from the 5-HT score list formed a statistically significant connected network (Fig. 6I, J). The set of genes connected in this network was enriched with Rho, p38 and non-canonical Wnt signaling pathways (Fig. 6J). Among main hubs of the network we found such an important for the process of development genes as cdc42, rac1, egr1, mapk14 and ctnnb1. Similar analysis using HPRD protein interaction network [46] also highlighted guanine-binding, RHOC-related and b-catenin-related subnetworks (Fig. S7).

## Discussion

The enigma of pre-neural serotonin and its role during early stages of embryonic development in different invertebrate animals inspired us to investigate its role in early pre-neural vertebrate development. Our previous work on the invertebrate model of *Lymnaea stagnalis* revealed that increase of 5-HT exclusively during early pre-neural developmental stages affects later developmental aspects and behavior of juvenile snails [38]. This study clearly suggested that mother-derived serotonin affects the lifestyle of progeny in a delayed and non-genetic manner. To investigate whether this holds true for vertebrates as well, we used zebrafish embryos and larvae to address the role of pre-neural serotonin. We found that increase rather than decrease of 5-HT level in zygotes and early cleaving embryos results in conceptually-similar delayed effects in juvenile and adult individuals.

We firstly analyzed the development of serotonergic system starting from a zygote stage. Consistently with earlier studies, we detected the presence of serotonin [21,22] as well as some receptors and AADC in cleaving embryos of teleost fish [23,47]. Our experiments demonstrated that pre-neural zebrafish embryos synthesize and capture 5-HT from the environment, and that this 5-HT is anisotropically distributed, being mostly present in a cleaving part of an embryo. Some earlier studies suggested that differential distribution of 5-HT in cleaving blastomeres may play the role of an instructive morphogenetic gradient [48]. However, unlike the situation in *Xenopus* [18], 5-HT did show any signs of a gradient in blastomeres of zebrafish. Moreover, when we experimentally created such a gradient, no major developmental processes were affected.

The presence of a multicomponent and complex serotonergic system at early pre-neural developmental stages suggested the importance of serotonin for key developmental transitions in fish. However, the pharmacological screen with a number of serotonergic system-related drugs did not reveal any disturbances in cleavage, gastrulation and other developmental processes, even in those that were previously reported to be affected in similar experimental settings in annelid worms [49], mollusks [14,16,38], insects [13], sea urchins [15], ascidians [50], amphibians [51] and mammals [52,53]. Importantly, we did not detect any abnormalities in left-right asymmetry development, which has been shown to be the case when serotonergic system was affected in amphibians and birds [17,18]. Overall, the early development of teleost fish turned out to be surprisingly stable and quite insensitive to the modulation of early serotonergic system. At the same time, multiple components of serotonergic system are present at early developmental stages of a fish and are conservative, rendering this aspect as highly similar to other vertebrate and invertebrate groups of animals. Thus, the pre-neural early developmental serotonin might play some other subtle roles, for instance, controlling delayed or mild effects operating in mid-, late- and post-embryonic development. To address such delayed effects of serotonin, we performed experiments with modulations of pre-neural embryonic 5-HT followed by the analysis of larval serotonergic system together with behavioral tests, morphology of larval body and measurements of monoamines and their metabolites. As a result, we did not detect any significant morphologic abnormalities in fish larvae that developed from the early-manipulated embryos. Unexpectedly, juvenile fish with modulated early pre-neural serotonergic system demonstrated subtle but significant changes in their behavior. These effects were restricted to the early pre-neural stage treatments, since disturbances of serotonergic system at any later time point, including stages of neural development, could not recapitulate the early stage-treatment effects. Thus, the early stage effects cannot be based on retained and stored serotonin. In nature, such mechanisms may play a potent adaptive role. For instance, the serotonin from the mother or from the immediate environment, where embryos develop, may define the fine aspects of young fish lifestyle.

These results reveal different roles of pre-neural and neural serotonin and suggest a new explanatory framework that clarifies some logical incongruence found in previously published data. For instance, previous studies demonstrated that experimental reduction of 5-HT achieved by blocking the TPH activity, or, alternatively, performing a gene knock out or inducing a pharmacological inhibition of AADC at 1-4 dpf, suppress fish movements with minor further recovery [29,47]. On the other hand, the elevation of 5-HT in fish results in some paradoxical effects. For instance, direct injection of 5-HT into the pericardium of 4 dpf larva leads to the increased locomotor activity [54], whereas the increase of 5-HT levels by inhibition of MAO from zygote to 5-7 dpf results in hypolocomotion of zebrafish larvae [55]. Furthermore, Sallinen et al. revealed that both general level of 5-HT and local content in specific tissues in such fish are significantly augmented. This hyperserotonic phenotype in larvae after MAO inhibition included noticeable decrease of 5-HT immunoreactivity in the serotonergic cell soma in the brain [55]. In our study, we found the similar reduction of anti-5-HT staining at 2 dpf achieved by the elevation of 5-HT level starting earlier at zygote stage. Another type of hyperserotonic phenotype characterized by hypolocomotion is observed in case of blocking SERT, and the decrease in swimming activity appears in case of incubation in SSRI only starting from 3 dpf. At the same time, drug administration between 10 and 72 hpf causes no alteration in locomotion [56]. Hence, all these contradictions can be explained by our concept of different action and role played by early pre-neural versus late serotonin during fish development. Indeed, the gene expression analysis showed that injection of 5-HT in zygote causes wide changes that show a trend coherent with our findings showing diverging roles of serotonin at different developmental stages. Numerous genes that should be switched on at 2dpf stayed downregulated and instead were seen upregulated at 4 dpf, when they should be already downregulated as compared to the stage-matched controls. Therefore, increased zygotic serotonin delays the expression of many genes, and at the same time it also causes the premature expression of some rather late genes. In the latter case, the number of such early upregulated genes is much lower as compared to the number of genes that are delayed in their expression level.

Complementary to our reasoning in the previous paragraph, it is worth mentioning that the catecholaminergic system is connected with the serotonergic system in the brain. In agreement with this, both pre-neural and early neural increase of 5-HT in our experiments resulted in upregulated catecholamine’s metabolism. The experiments involving MAO inhibition did not affect the content of catecholamines themselves, but led to the decrease of MAO-related dopamine metabolite DOPAC [55]. Comparisons with data from the literature revealed that MAO blocking described by Sallinen et al. and application of 5-HT or 5-HTP in our experiments caused similar order of changes in 5-HT concentrations. Despite this, the effects on larval behavior were much less pronounced in case of 5-HT or 5-HTP administration. Thus, hyperserotonic phenotype can develop also via influencing catecholaminergic system in addition to serotonergic system-mediated effects.

In our experiments, in addition to particular changes in larval behavior in response to the early serotonin manipulation, we observed that serotonergic neurons in 2 dpf embryos showed significantly less serotonin after 5-HT application to zygote or early cleavage stages (but not to the later stages of development). Serotonergic neurons in the dorsal root nuclei mediate short-term motor learning in zebrafish larva [57]. These neurons activation decreases acoustic startle response habituation while 5-HT depletion leads to its enhancing [58]. Similarly to Sallinen with co-authors [55], we could clearly see the reduction of 5-HT in all types of serotonergic cells as a part of hyperserotonic phenotype. This invariably implies that 5-HT synthesis is affected in all cells where it depends on different isoforms of TPH. At the same time, the transcriptomics profiling did not reveal significant shifts in the expression of enzymes and transporters related to 5-HT metabolism and turnover, which suggests an alternative molecular mechanism.

One of the possible explanations might include different mechanisms of a post-translational control of TPH activity. In *Sert^-/-^* mice, the *in vivo* level of 5-HT synthesis is upregulated without changes in expression and *in vitro* activity of both TPH isoforms [59]. The mechanisms of such control can include phosphorylation of TPH, synthesis and regeneration of tetrahydrobiopterin coenzyme, changes in iron metabolism or inactivation of TPH by reactive oxygen species [60]. Unexpectedly, 5-HT and its metabolite 5-HIAA showed different levels in the head and the rest of the body of advanced larvae after 5-HT administration into a zygote. Therefore, we observed a conceptual difference between metabolism of 5-HT in a head and in the rest of the body in response to the early pre-neural increase of 5-HT concentration. The subtle difference between the results of anti-5-HT IHC staining and direct measurements based on HPLC might be explained by masking of intracellular decrease of 5-HT with increased level of 5-HT in blood and intercellular spaces. Additionally, in the brain, local 5-HT synthesis and degradation might be regulated by inhibitory neuronal feedback loops [61]. Our results suggest that such regulation may exists not only within neurons but also in other cell types at different time points during development.

The delayed nature of effects of pre-neural serotonin is not based on the retained serotonin since we tested for this by performing later treatments. On the other hand, the observed phenomenon requires an explanation at the molecular level. In principle, the delayed effects can be transmitted through the intermediate developmental stages via the states of secondary messenger systems, retained expression of transcription factors, changes in epigenetic landscapes or post-translational modifications of proteins including their serotonylation (post-translational control of a protein function, which is based on the covalent binding of serotonin to the residues of glutamines). Serotonylation directly depends on the intracellular concentration of serotonin, and, thus, can be a convenient mean of delaying and relaying a signal in our settings. The potential role of serotonylation in 5-HT-related aspects of vertebrates development can be similar to its signal-relaying function revealed in our previous study on *Lymnaea stagnalis* [38]. However, our experiments did not support its direct involvement into the early 5-HT-related regulations in zebrafish. The reason of this may be in fact that during zebrafish development unlike in *Lymnaea stagnalis* or mammals, the oocyte size is large while the volume of the body does not increase much between a zygote and a first feeding stage. This means that 5-HT deposited into an egg by the mother or 5-HT captured from the environment would be retained, unless new synthesis or intense catabolism is involved. This might enable a fine 5-HT-dependent control at any pre-neural or neural developmental step, including delayed effects in the presence of a strong catabolism or later body growth dissolving the original concentration. Once deposited by a mother, the early pre-neural serotonin can cause the delayed effects such as affecting the formation of different populations of differentiating neurons, which, in turn, might “imprint” specific behavioral features retained for the rest of the animal’s life (Fig. 6L). In line with this, our results demonstrate that the 5-HT content in a zygote and synthesis of 5-HT in the early embryonic neurons are interlinked by a negative feedback loop. This is fully consistent with the fact that early pre-neural elevation of 5-HT causes an increase of a short-term behavioral habituation.

Thus, natural variation of 5-HT level in a zygote may represent one of the sources for the inter-individual behavioral plasticity. In our previous work on *L. stagnalis*, we found that 5-HT is released from the embryo to the environment [38]. Capacity of pre-neural zebrafish embryos to uptake 5-HT from the media suggests that 5-HT may additionally serve as a quorum sensing molecule mediating communication between developing embryos as well as for sensing the environment in general, which might be necessary for the formation of the adaptive behavioral diversity in juveniles and adult fish (Fig. 6L).

## Materials and Methods

### Animal maintenance and general manipulation procedures

For experiments we used AB line fish kept in Karolinska Institutet zebrafish core facility. All works were performed according to Swedish laws and regulations on animal experimentation. Fish eggs were obtained by natural fertilization process with the standard procedures described before [62,63].

### Pharmacological treatments

Injections of embryos were performed using Eppendorf FemtoJet microinjector. We co-injected substances with lysine-fixable TRITC-conjugated fluorescent dextran (Molecular Probes, USA). To avoid impact of differences in permeability, we dechorionated zebrafish eggs at zygote stage using proteinase K protocol [62] for the most of the pharmacological treatments.

The following drugs were used for pharmacological treatment (produced by Tocris, unless otherwise specified): serotonin (5-hydroxytryptamine hydrochloride, 5-HT, Sigma-Aldrich); L-tryptophan (tryptophan, Trp, Sigma-Aldrich); 5-hydroxy-L-tryptophan (5-HTP, Sigma-Aldrich); 3-hydroxybenzilhydrazine (NSD-1015, Sigma-Aldrich); m-CPP hydrochloride (mCPP); 4-iodo-2,5-dimethoxy-α-methylbenzene ethanamine hydrochloride (DOI); mianserin hydrochloride (mianserin); spiperone hydrochloride (spiperone); ritanserin tartrate (ritanserin); methysergide maleate (methysergid); cyproheptadine hydrochloride (cyproheptadine); trazodone hydrochloride (trazodone, Sigma-Aldrich); ketanserin tartrate (ketanserin), citalopram hydrobromide (citalopram), clomipramine hydrochloride (clomipramine), fluoxetine hydrochloride (fluoxetine), fluvoxamine hydrochloride (fluvoxamine), imipramine hydrochloride (imipramine), cystamine dihydrochloride (cystamine, Sigma-Aldrich), monodansylcadaverin (MDC, Sigma-Aldrich), 5-carboxamidotryptamine (5-CT), 5-methoxy-N,N-dimethyltryptamine (5-MeO-DMT), 7-(Dipropylamino)-5,6,7,8-tetrahydronaphthalen-1-ol (8-OH-DPAT), α-Methylserotonin (αMS), (6aR,9R)-5-bromo-N,Ndiethyl-7-methyl-4,6,6a,7,8,9-hexahydroindolo[4,3-fg]quinoline-9-carboxamide (Br-LSD, kindly gift of Dr. Laszlo Hiripi), tropanyl-3,5-dimethylbenzoate (tropanyl), (S)-WAY 100135 dihydrochloride (WAY-100.135), SB 269970 hydrochloride (SB269970), GR-113808, zacopride hydrochloride (zacopride), NAN-190 hydrobromide (NAN-190) all mineral salts were obtained from Sigma-Aldrich. 5-HT, 5-HTP and tryptophan solutions of specified concentrations were freshly prepared in embryo medium. All other solutions were prepared as 10 mM stocks in DMSO, d.i. water of 95% ethanol and stored at −20°C prior to application to the embryo medium.

### Immunohistochemistry, image acquisition and analysis

For IHC staining dechorionated embryos and larvae were fixed overnight in 4% PFA on 1x PBS. Embryos and larvae up to 2 dpf were permeabilized with 5% triton X-100 solution in PBS (overnight, 4°C). 4 dpf fish were additionally treated with proteinase K [62]. Samples were incubated in polyclonal rabbit anti-5-HT antibody (ImmunoStar, Hudson, USA, #20080, polyclonal, rabbit, dilution 1:2000) during 48-72 hours at 10°C. Primary antibodies were detected with donkey-anti-rabbit Alexa 488 and Alexa 633 -conjugated IgG (Molecular Probes, USA, diluted 1:800 in PBS), 8 h at 4°C. Cell nuclei were stained with DAPI or Hoechst 33342. After 3×10 min washing in PBS, the specimens were immersed in 2,2’-thiodiethanol (TDE) and placed to the glass bottom dishes for microscopic analysis and image acquisition.

For 5-HT level estimation, we obtained confocal images of immunostained early embryos. Scans were performed with fully open pinhole. Measurements and analysis were performed with FIJI software. Entire embryos or relevant parts were manually delineated, and average pixel intensity was obtained inside of the border in arbitrary fluorescence units (a.u.). Measurements of 5-HT level in the brain cells were done in volume basing on the full 3D scans of fish head. We manually traced each slice for the segmentation of cell volumes using Imaris software (Bitplane), then estimated mean brightness within volumes delimited by the 3D surfaces.

### Measurements of monoamines and their metabolites

Whole fish or freshly dissected heads were homogenized at 0°C with 100 μl of 0.2 M perchloric acid containing 100 μM EDTA-2Na. Following standing for 30 min on ice, the homogenates are centrifuged for 5 min at 12,000 x g at 4°C. The supernatants are carefully aspirated and mixed with 1M sodium acetate buffer at a ratio 5:1 (v/v). Resulting solution was filtered through a centrifuge filter (30K PES membrane, VWR) for 7 min at 12,000 x g at 4°C. The filtrates were analyzed immediately.

Measurements of monoamines concentrations were performed using high pressure liquid chromatography (HPLC) with electrochemical detection as described elsewhere [64]. Briefly, the HPLC system consisted of a HTEC500 unit (Eicom, Kyoto, Japan), and a CMA/200 Refrigerated Microsampler (CMA Microdialysis, Stockholm, Sweden) equipped with a 20 μL loop and operating at +4°C. The potential of the glassy carbon working electrode was +450 mV versus the Ag/AgCl reference electrode. The separation was achieved on a 200 × 2.0 mm Eicompak CAX column (Eicom). The mobile phase was a mixture of methanol and 0.1 M phosphate buffer (pH 6.0) (30:70, v/v) containing 20 mM potassium chloride and 0.13 mM EDTA-2Na. The chromatograms were recorded and integrated by use of a computerized data acquisition system Clarity (DataApex, Prague, Czech Republic). The detection limit (signal-to-noise ratio = 3) for NA, DA and 5-HT was 0.5 fmol in 10 μl injected onto the column respectively.

Concentrations of DOPAC, HVA and 5-HIAA were determined by a separate HPLC system with electrochemical detection (HTEC500). The potential of the glassy carbon working electrode was +750 mV versus the Ag/AgCl reference electrode. The separation was achieved on a 150 × 3.0 mm Eicompak SC-5ODS column (Eicom). The mobile phase was a mixture of methanol and 0.1 M citrate/0.1 M sodium acetate buffer solution (pH 3.5) (16:84, v/v) and contained 0.971 mM octanesulphonic acid sodium salt and 0.013 mM EDTA-2Na. The detection limit (signal-to-noise ratio = 3) for DOPAC, HVA and 5-HIAA was 20 fmol in 10 μl injected onto the column respectively. The chromatograms were recorded and integrated by use of the computerized data acquisition system Clarity (DataApex).

In order to determine number of larvae per sample essential for measurements, we built normalization dependences of concentration to fish number which showed linear character and thus concentrations were expressed in quantity of substance per fish or fish’s head.

### Behavioral tests

For the analysis of spontaneous coiling activity in 1 dpf larvae we recorded 5 min footages, then manually counted the number of contraction. The response to touch was analyzed in 2-4 dpf larvae. Each larva was repeatedly stimulated by gently touching it with a needle in the trunk region. A significant response was recorded when the stimulation was followed by a displacement of at least 1 body length. Each fish was stimulated ^~^50 times (^~^1 touch per second). All experiments were videorecorded and analyzed manually.

Behavioral assays on 5 and 6 dpf fish were performed using integrated high throughput systems for videotracking and environmental control (DanioVision, Noldus, Wageningen, Netherlands) as previously described [63]. The larvae were plated individually in 48-well plates (round wells, 10 mm in diameter) at least 1h before testing. All experiments were performed during the light phase of the diurnal cycle (light/dark 14h/10h). Spontaneous swimming was recorded under white light illumination for 5 min before the dark pulse. A sound stimulus was applied by means of a solenoid plunger hitting the base of the recording chamber. Both white light illumination and acoustic stimulation were controlled by the videotracking software and synchronized with the video recording. Swimming activity was recorded under constant infrared illumination at 50 fps and tracked in live mode. The XY coordinates were exported as ASCII files and analyzed using custom-made routines implemented in Matlab™ (The Mathworks, Nattick, MD, USA).

The acoustic startle response was first evaluated by averaging the response to 10 consecutive stimulations applied with 1 min intervals to avoid habituation. The habituation to acoustic stimulation was assessed by decreasing the interval between stimuli to 10 s.

The dark pulse experiment consisted of a 10-min long interval when the light was turned off during the light phase of the dark/light cycle. Swimming activity was tracked for 10 min before, during the dark episode, and for 10 min after the turning the light back on. We analyzed the dark pulse-induced hyperactivity, the gradual decline in activity level during the dark pulse and the response to turning the light back.

### PCR and qPCR

40-60 fish embryos or larvae were placed into trizol reagent and total RNA was extracted using the RNeasy Mini Kit (Qiagen). 1 μg of total RNA was treated with RQ1 Rnase-free DNase (Promega) and reverse transcribed using SuperScript II Reverse Transcriptase (Invitrogen) and random primers (RT+ reaction). Parallel reactions without reverse transcriptase enzyme were performed as a control (RT–reaction), and SYBR Green I real-time quantitative PCR assays were carried out, as described in [65]. Expression levels were obtained by normalization to the value of housekeeping genes, obtained for every sample in parallel assays. For the information about the primers used to obtain the results presented on Fig.1I and 5L-M, see the Supplementary Tables 2 and 3 respectively.

### Microarray assay

Total RNA was extracted from 30-40 2 and 4 dpf zebrafish larvae from each experimental group. Total RNA from each sample was used for the synthesis of cDNA, which was hybridized to 902007 Affymetrix® Zebrafish Gene 1.0 ST Array with Ambion WT terminal labeling and hybridization protocol. In the annotation provided by Affymetric for Zebrafish Gene 1.0 ST Array chip, 24497 probesets from 73244 were annotated by gene names. We re-mapped the sequences of the probesets to Zv9/danRer7 reference genome of zebrafish and used the RefSeq mRNA annotation track from UCSC genome browser annotation database to assign names for 30784 probesets in total, which were used in further analysis.

Two replicates showed good correlation between both controls and 5-HT-induced conditions (Pearson correlation coefficient calculated for all probesets, r=0.98, p-value<10^−16^). Therefore, in the calculations we always used average values between two replicates.

Gene Set Enrichment Analysis (GSEA) was performed using the standard GSEA software, pre-ranked analysis [66] was done with Gene Ontologies for zebrafish. The gene ranks were formed by selecting the probesets showing the maximal by absolute value difference between two compared conditions. All probesets annotated by gene names have been considered in this analysis, which resulted in the values associated to 20691 distinct gene names.

The transcriptomic data discussed in this publication have been deposited in NCBI’s Gene Expression Omnibus and are accessible through GEO Series accession number GSE122201.

### Statistical analysis

For the statistical analysis we used Statistica 10 (StatSoft), XLstat (Addinsoft) or Prism 7 (GraphPad) software packages. All data are presented as average ± s.e.m. or 95% confidence interval (see Figure legends). Unpaired Student’s t-test was used when two groups were compared after the normal distribution test was passed otherwise ANOVA test was applied. A non-linear regression with one phase exponential decay equation and further comparison of the equation parameters with an extra sum-of-square F-test was used to estimate differences in the habituation to startle response. More detailed information on the statistical analysis is given in the Figure legends. Differences were taken as statistically significant in case of p<0.05.

## Supporting information

Supplementary File 1. Microarray Data

Supplementary File 2. Tables 1 & 2, PCR Primers Sequences

---

**Fig. S1.**
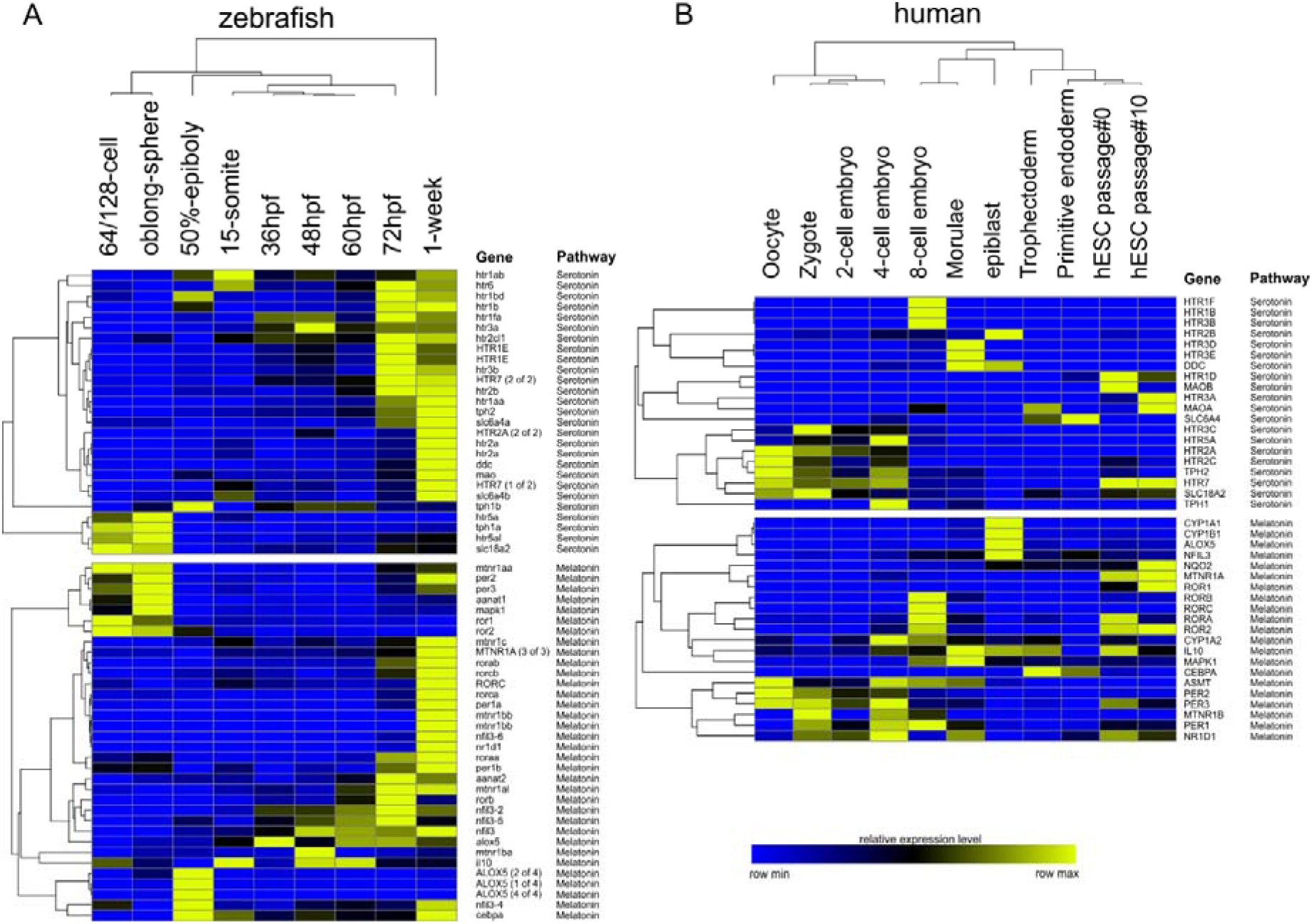
Expression of 5-HT-related, melatonin-related and kynurenine-related genes during early development of zebrafish and human. Hierarchical clustering of developmental transcriptomics data for zebrafish (A) and human (B). Data were obtained from public sources (see the text).

**Fig. S2.**
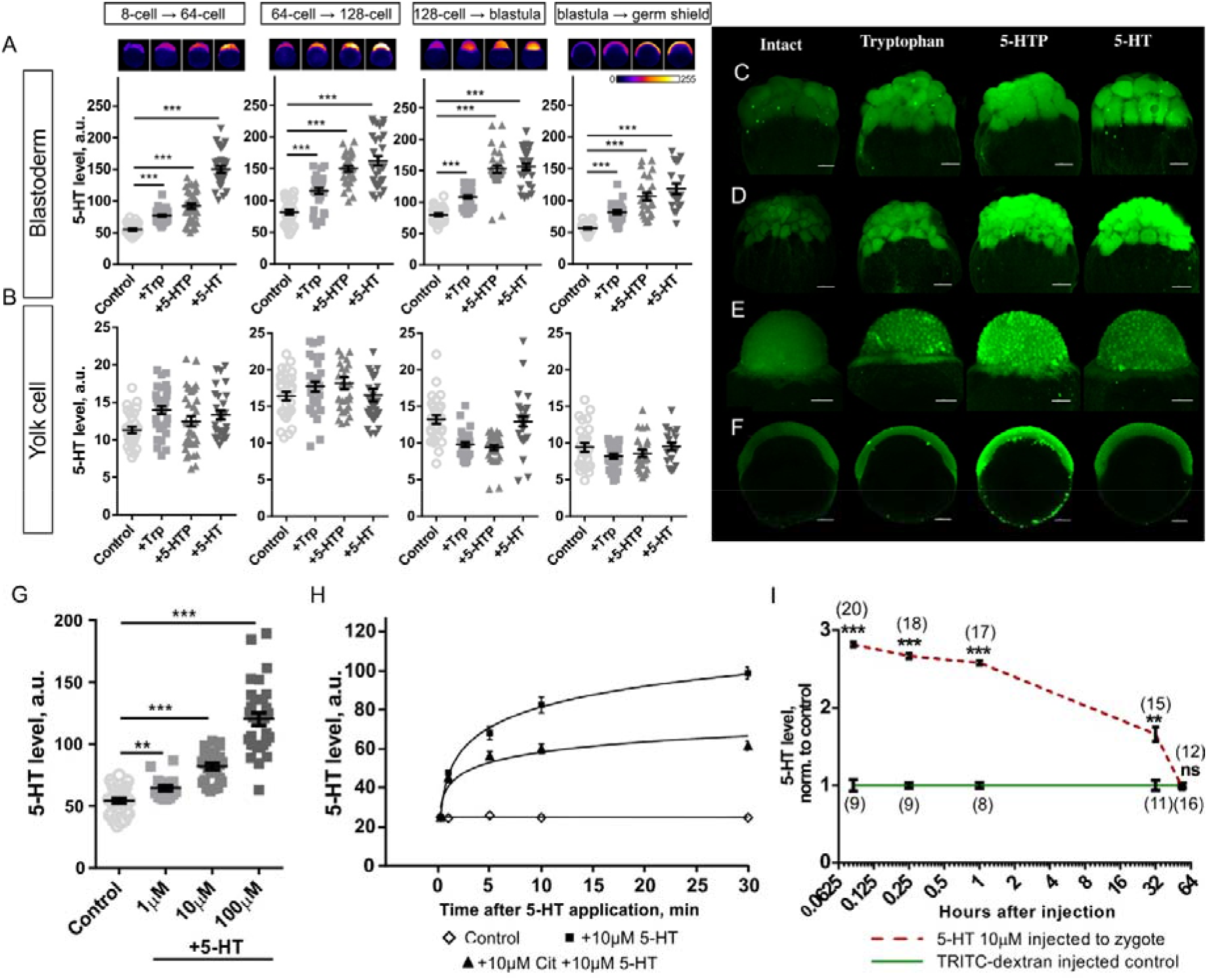
Synthesis and transport of 5-HT in an early zebrafish embryo. (A-B) – 5-HT-levels measured as a relative brightness of a 5-HT immunostaining in the cleaving part (A) and the yolk cell (B) of an embryo after incubations in tryptophan, 5-HTP and 5-HT at different stages of zebrafish early development. (C-F) – 5-HT-immunostaining of embryos after incubations in tryptophan, 5-HTP and 5-HT. Incubation intervals are the same as shown in (A-B). (G-H) – 5-HT-levels (relative brightness of a 5-HT immunostaining) in the cleaving part of an embryo. (G) - 5-HT was applied in different concentrations for 1 hour at a 2-cell stage. (H) – change of 5-HT-level through the time after the application of 5-HT at 2-cell stage with or without SSRI citalopram. (I) – Change of 5-HT-level (measured as a relative brightness of a 5-HT immunostaining) over time in the whole embryo volume after the injection into a zygote, folds change to control. The sample size for each point is given in parentheses. All data represented as mean ± s.e.m.; T-test; ns – non-significant; ** – p<0.01; *** – p<0.001. Scale bars: (C-F) – 100 μm.

**Fig. S3.**
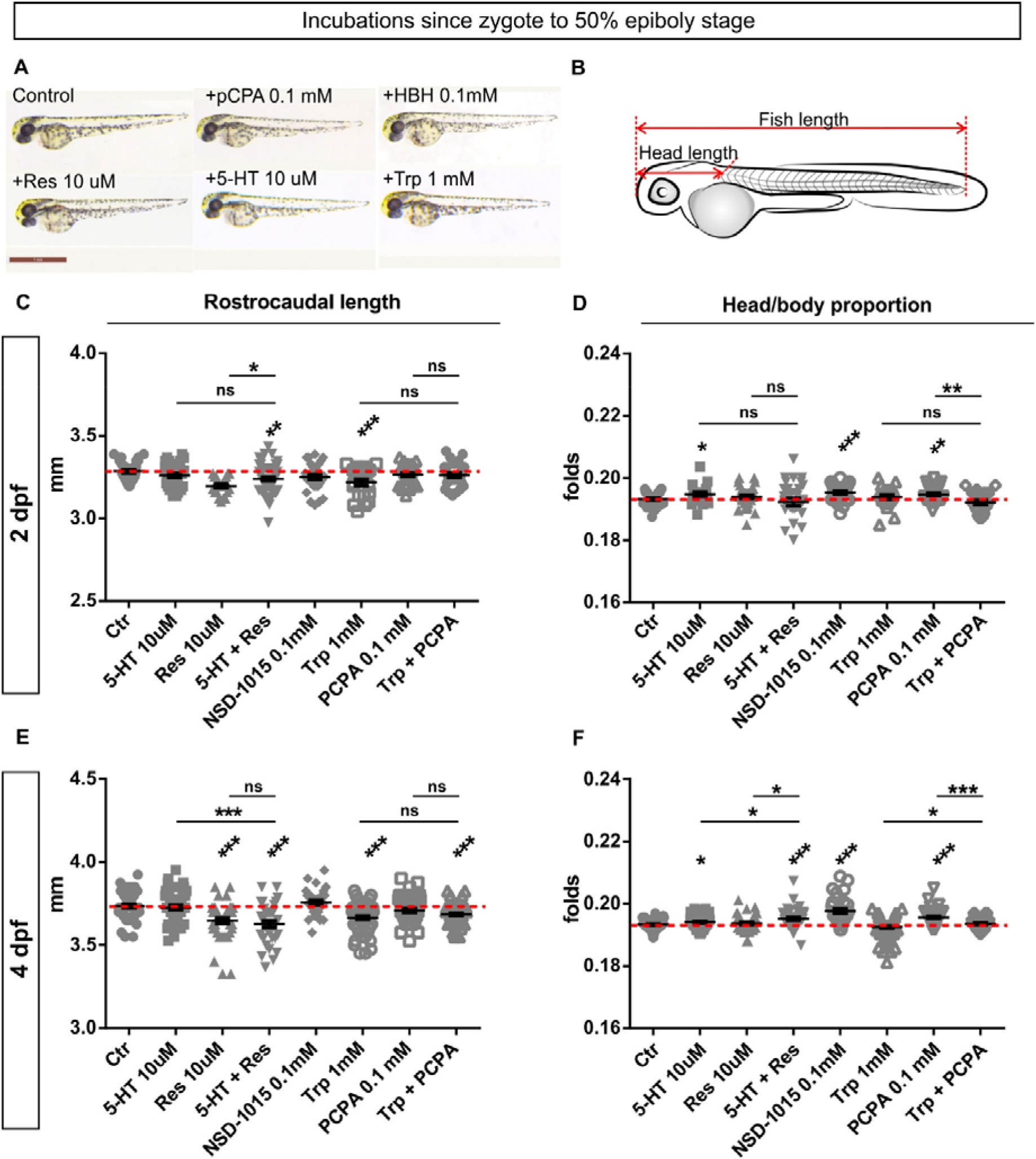
Rostrocaudal length in zebrafish embryos after the exposure to 5-HT and 5-HT-related substances during early pre-neural developmental stages. (A) – Examples of representative 2 dpf fish larvae after early pre-neural application of 5-HT, it’s precursors and blockers of synthesis. (B) – Scheme of measurements. (C-E) – Comparison of rostrocaudal fish length (C, E) and head/body proportions (D, F) in 2 dpf (C, D) and 4 dpf (E, F) larvae. Data represented as mean ± s.e.m.; n = 20-110; T-test; * – p<0.05; ** – p<0.01; *** – p<0.001.

**Fig. S4.**
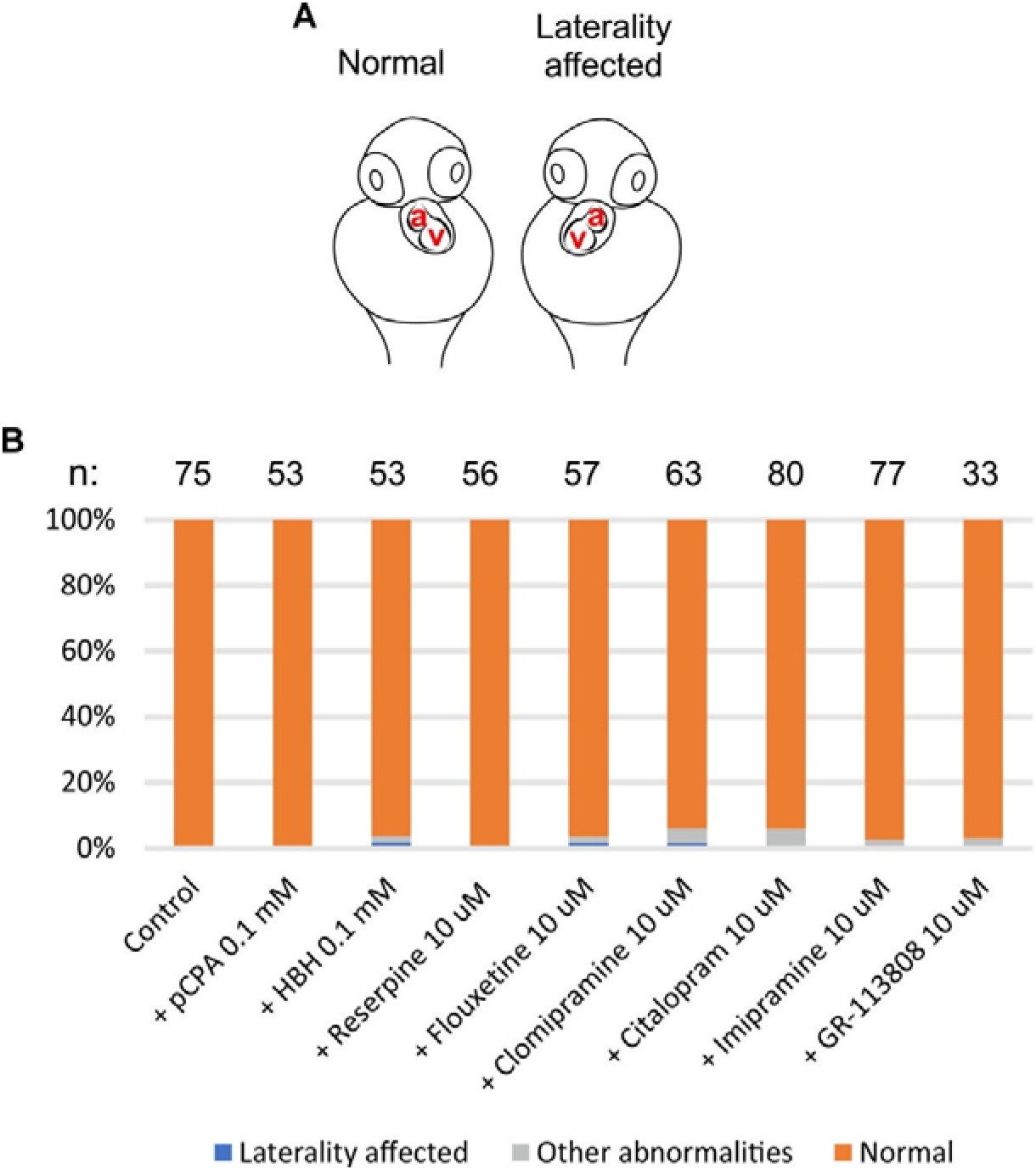
Left-right polarity is not affected after the exposure of cleaving zebrafish embryos to the selective 5-HT_4_ receptor antagonist GR-113808 and inhibitors of a 5-HT synthesis and transport. (A) – Left-right polarity abnormalities of the heart. Atrioventricular heart asymmetry orientation in 2-3 dpf larvae was analyzed. (B) – Ratio of heart asymmetry abnormalities in embryos exposed to different substances starting from a zygote until the late blastula stage.

**Fig. S5.**
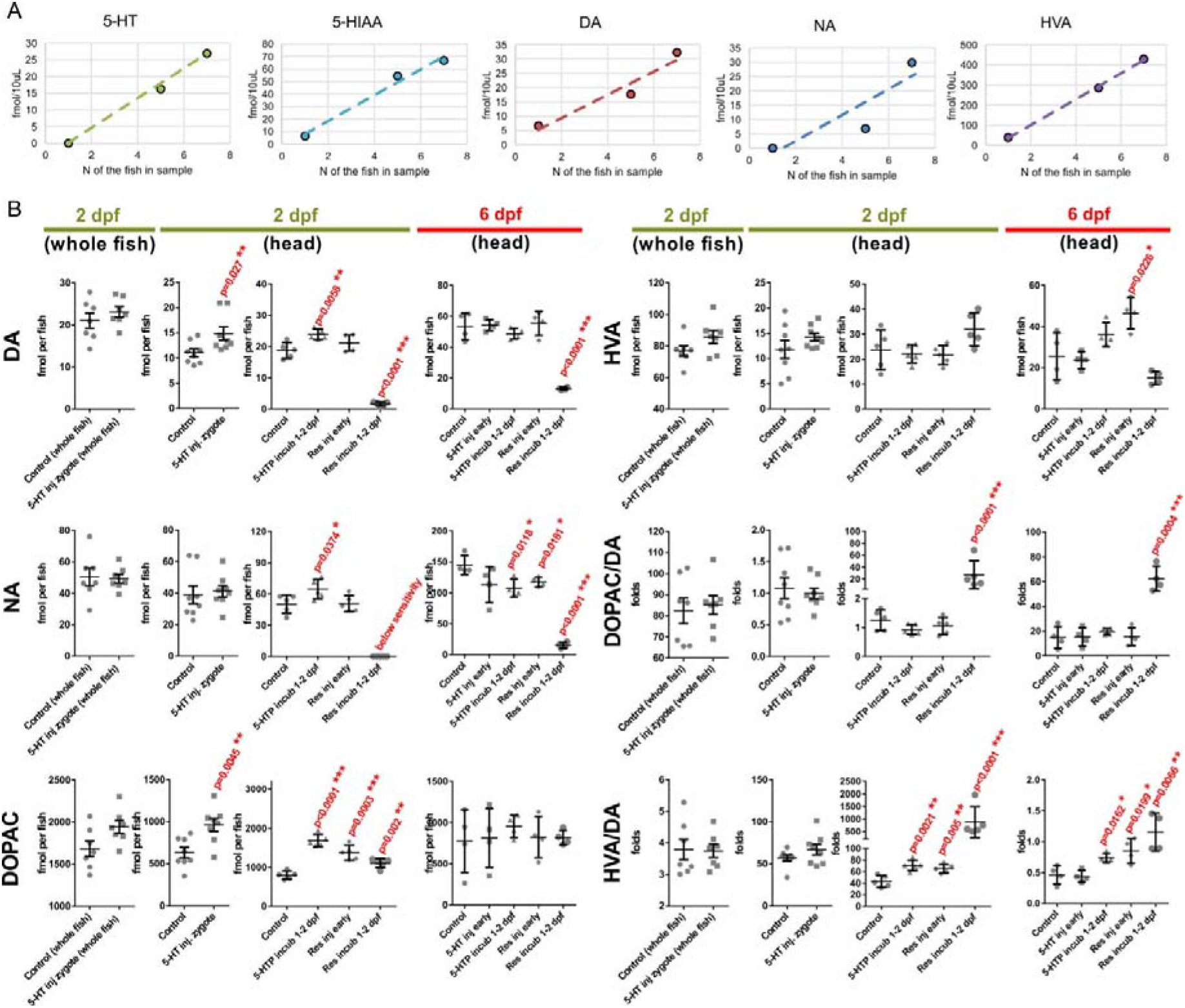
HPLC-measurements of monoamines and their metabolites in fish after modulations of 5-HT levels at pre-neural (zygote) and early neural (1-2 dpf) developmental stages. (A) – Calibration curves for the measured substances (experiment performed with different numbers of 2-dpf larvae per measured sample). (B) – Levels of dopamine (DA), noradrenaline (NA), dopamine metabolites 3,4-dihydroxyphenylacetic acid (DOPAC) and homovanillic acid (HVA) in the whole 2 dpf zebrafish larvae and in the severed heads of 2 dpf and 6 dpf fish. Experimental groups and conditions are the same as shown in the Fig. 3(A-D).

**Fig. S6.**
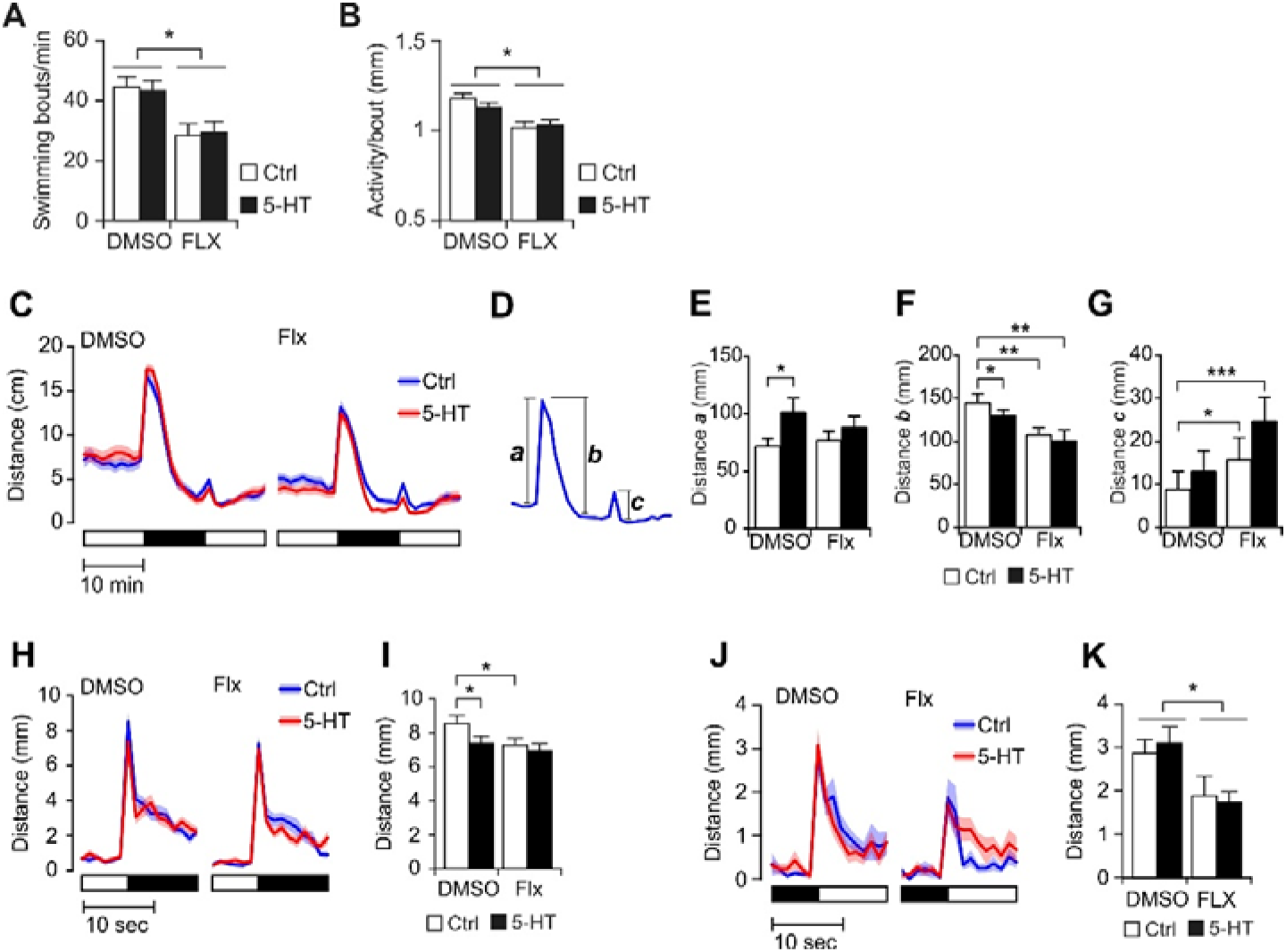
Analysis of 5 dpf larvae behavior after the early 5-HT increase and 4-5 dpf exposure to fluoxetine. (A-C) – Spontaneous swimming activity. (A) – Speed, (B) – bouts frequency, (C) – displacement per bout. (D-H) – Results of the dark pulse test for 5 dpf fish. The larvae were subjected to a 10-min interval of complete darkness. Swimming activity was recorded continuously starting 10 min before the dark pulse, in darkness and for 10 min after the end of the dark pulse. Experimental groups were the same as described in Fig 4D. (I-L) – Hyperactivity in response to sudden change in light intensity. (I, J) – Turning the light off triggers a stress response and is followed by sustained increase in activity. 5-HT injection at zygote stage, or Flx incubation for 24h prior to testing decrease the amplitude of the response. (K, L) – Turning the light on (virtually instantaneous increase in white light intensity from 0 to ^~^300 lux) triggers a startle response followed by transient increase in spontaneous activity. The amplitude of the startle response is reduced by incubation with Flx for 24h prior to testing. All data represented as mean ± s.e.m.; factorial ANOVA followed by student’s t-test for pair-wise comparisons; n=150-200 larvae; * – p<0.05; ** – p<0.01.

**Fig. S7.**
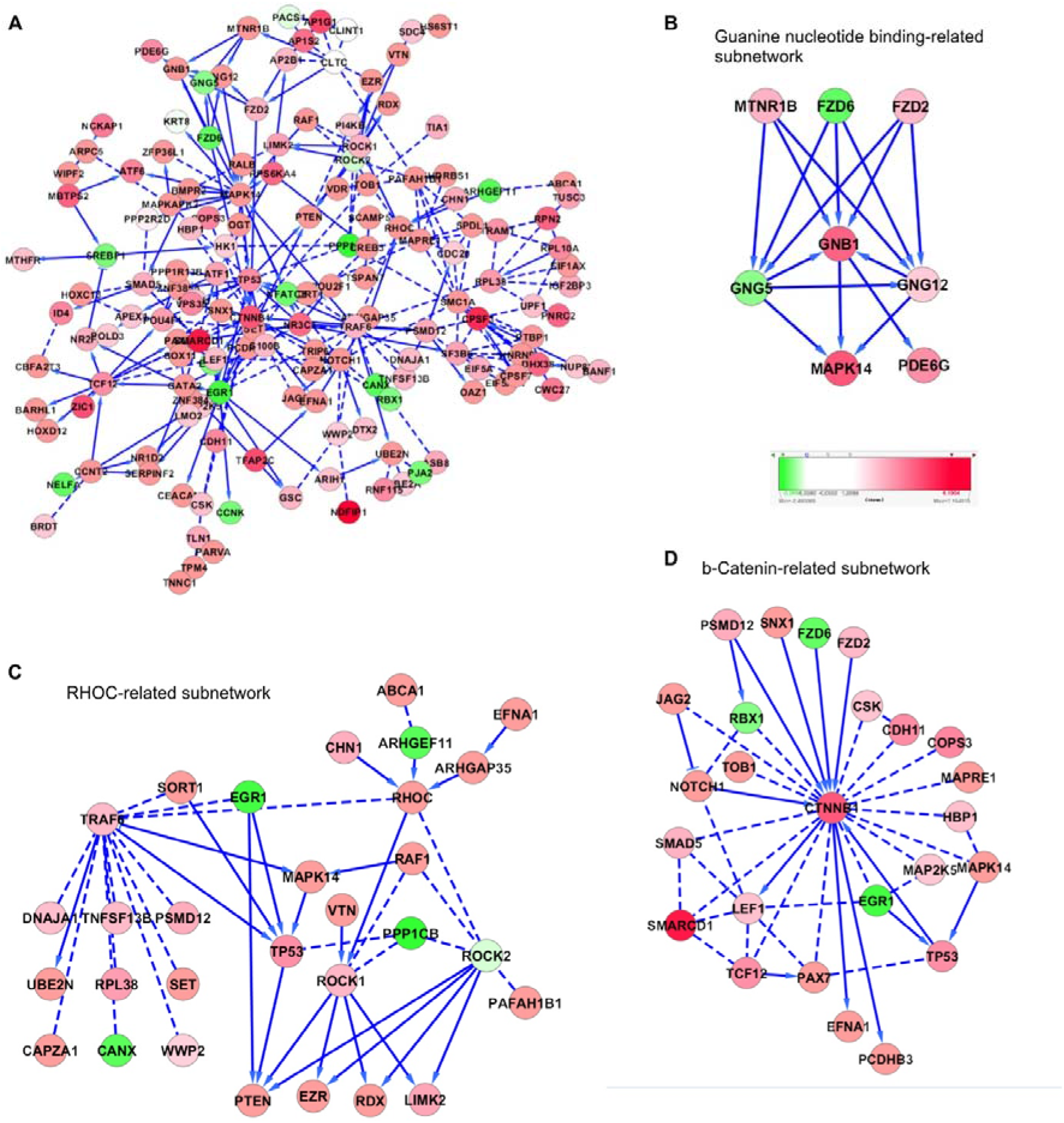
Reactome functional interaction network analysis with OFTEN method. (A) – The largest connected component formed by functional interactions from REACTOME database between 400 top scored 5-HT-effect genes, with both signs, positive (shown by red) and negative (shown by green). Dashed lines correspond to functional interactions of low confidence (mostly, predicted by computational methods or deduced from co-participation in a molecular complex). (B) – Guanine nucleotide binding-related subnetwork (all proteins functionally interacting with GNB1 through one interaction). (C) – RHOC-related subnetwork (all proteins functionally interacting with RHOC through one or two interactions). (D) – β-catenin-related subnetwork (all proteins functionally interacting with CTNNB1 through one interaction).

Supplementary File 1: Transcriptomic data collected for assessing the transcriptomic response to injections of 5-HT and reserpine to zygote at day 2 and day 4 of the developmental course (Excel file). Results of the gene set enrichment analysis (GSEA). List of anatomical GO terms obtained using ZEOGS tool.

Supplementary File 2: PCR primers used in this study.

## Notes

### Competing Interest Statement

The authors have declared no competing interest.

## Bibliography

1. Sloley BD (2004) Metabolism of monoamines in invertebrates: the relative importance of monoamine oxidase in different phyla. Neurotoxicology 25: 175–183. doi:10.1016/S0161-813X(03)00096-2.

2. Berger M, Gray JA, Roth BL (2009) The expanded biology of serotonin. Annu Rev Med 60: 355–366. doi: 10.1146/annurev.med.60.042307.110802.

3. Bianchi P, Pimentel DR, Murphy MP, Colucci WS, Parini A (2005) A new hypertrophic mechanism of serotonin in cardiac myocytes: receptor-independent ROS generation. FASEB J 19: 641–643. doi:10.1096/fj.04-2518fje.

4. Ren G, Li S, Zhong H, Lin S (2013) Zebrafish tyrosine hydroxylase 2 gene encodes tryptophan hydroxylase. J Biol Chem 288: 22451–22459. doi:10.1074/jbc.M113.485227.

5. Ramamoorthy S, Samuvel DJ, Buck ER, Rudnick G, Jayanthi LD (2007) Phosphorylation of threonine residue 276 is required for acute regulation of serotonin transporter by cyclic GMP. J Biol Chem 282: 11639–11647. doi:10.1074/jbc.M611353200.

6. Marin P, Chanrion B, Bockaert J (2007) Quand deux molécules impliquées dans la régulation de l’humeur se rencontrent. Med Sci (Paris) 23: 554–555. doi:10.1051/medsci/2007235554.

7. Carneiro AMD, Cook EH, Murphy DL, Blakely RD (2008) Interactions between integrin alphaIIbbeta3 and the serotonin transporter regulate serotonin transport and platelet aggregation in mice and humans. J Clin Invest 118: 1544–1552. doi:10.1172/JCI33374.

8. Muma NA, Mi Z (2015) Serotonylation and transamidation of other monoamines. ACS Chem Neurosci 6: 961–969. doi:10.1021/cn500329r.

9. Lillesaar C (2011) The serotonergic system in fish. J Chem Neuroanat 41: 294–308. doi:10.1016/j.jchemneu.2011.05.009.

10. Lawrence AJ, Soame JM (2009) The endocrine control of reproduction in Nereidae: a new multi-hormonal model with implications for their functional role in a changing environment. Philos Trans R Soc Lond B, Biol Sci 364: 3363–3376. doi:10.1098/rstb.2009.0127.

11. Stricker SA, Smythe TL (2000) Multiple triggers of oocyte maturation in nemertean worms: the roles of calcium and serotonin. J Exp Zool 287: 243–261. doi:10.1002/1097-010X(20000801)287:3<243::AID-JEZ6>3.0.CO;2-B.

12. Hanocq-Quertier J, Baltus E (1981) Induction of meiosis of Xenopus laevis oocytes by mianserine. Gamete Res 4: 49–56. doi:10.1002/mrd.1120040108.

13. Colas JF, Launay JM, Maroteaux L (1999) Maternal and zygotic control of serotonin biosynthesis are both necessary for Drosophila germband extension. Mech Dev 87: 67–76.

14. Buznikov GA, Nikitina LA, Voronezhskaya EE, Bezuglov VV, Dennis Willows AO, et al. (2003) Localization of serotonin and its possible role in early embryos of Tritonia diomedea(Mollusca: Nudibranchia). Cell Tissue Res 311: 259–266. doi:10.1007/s00441-002-0666-0.

15. Buznikov GA, Peterson RE, Nikitina LA, Bezuglov VV, Lauder JM (2005) The pre-nervous serotonergic system of developing sea urchin embryos and larvae: pharmacologic and immunocytochemical evidence. Neurochem Res 30: 825–837. doi:10.1007/s11064-005-6876-6.

16. Voronezhskaya EE, Khabarova MY, Nezlin LP, Ivashkin EG (2012) Delayed action of serotonin in molluscan development. Acta Biol Hung 63 Suppl 2: 210–216. doi:10.1556/ABiol.63.2012.Suppl.2.28.

17. Fukumoto T, Blakely R, Levin M (2005) Serotonin transporter function is an early step in left-right patterning in chick and frog embryos. Dev Neurosci 27: 349–363. doi:10.1159/000088451.

18. Fukumoto T, Kema IP, Levin M (2005) Serotonin signaling is a very early step in patterning of the left-right axis in chick and frog embryos. Curr Biol 15: 794–803. doi:10.1016/j.cub.2005.03.044.

19. Vandenberg LN, Lemire JM, Levin M (2013) Serotonin has early, cilia-independent roles in Xenopus left-right patterning. Dis Model Mech 6: 261–268. doi:10.1242/dmm.010256.

20. Beyer T, Danilchik M, Thumberger T, Vick P, Tisler M, et al. (2012) Serotonin signaling is required for Wnt-dependent GRP specification and leftward flow in Xenopus. Curr Biol 22: 33–39. doi:10.1016/j.cub.2011.11.027.

21. Bashammakh S, Seyfried S, Bader M, Kotnik Halavaty K (2014) Serotonin is required for pharyngeal arch morphogenesis in zebrafish. ScienceOpen Res. doi:10.14293/S2199-1006.1.SOR-LIFE.AWPDLZ.v1.

22. Buznikov GA, Chudakova IV, Zvezdina ND (1964) The role of neurohumours in early embryogenesis. i. serotonin content of developing embryos of sea urchin and loach. J Embryol Exp Morphol 12: 563–573.

23. Pei S, Liu L, Zhong Z, Wang H, Lin S, et al. (2016) Risk of prenatal depression and stress treatment: alteration on serotonin system of offspring through exposure to Fluoxetine. Sci Rep 6: 33822. doi:10.1038/srep33822.

24. Ori M, De Lucchini S, Marras G, Nardi I (2013) Unraveling new roles for serotonin receptor 2B in development: key findings from Xenopus. Int J Dev Biol 57: 707–714. doi:10.1387/ijdb.130204mo.

25. Bhasin N, LaMantia AS, Lauder JM (2004) Opposing regulation of cell proliferation by retinoic acid and the serotonin2B receptor in the mouse frontonasal mass. Anat Embryol (Berl) 208: 135–143. doi: 10.1007/s00429-004-0380-7.

26. Barreiro-Iglesias A, Mysiak KS, Scott AL, Reimer MM, Yang Y, et al. (2015) Serotonin promotes development and regeneration of spinal motor neurons in zebrafish. Cell Rep 13: 924–932. doi:10.1016/j.celrep.2015.09.050.

27. Doze VA, Perez DM (2012) G-protein-coupled receptors in adult neurogenesis. Pharmacol Rev 64: 645–675. doi:10.1124/pr.111.004762.

28. Pérez MR, Pellegrini E, Cano-Nicolau J, Gueguen M-M, Menouer-Le Guillou D, et al. (2013) Relationships between radial glial progenitors and 5-HT neurons in the paraventricular organ of adult zebrafish – potential effects of serotonin on adult neurogenesis. Eur J Neurosci 38: 3292–3301. doi:10.1111/ejn.12348.

29. Airhart MJ, Lee DH, Wilson TD, Miller BE, Miller MN, et al. (2012) Adverse effects of serotonin depletion in developing zebrafish. Neurotoxicol Teratol 34: 152–160. doi:10.1016/j.ntt.2011.08.008.

30. Prasad P, Ogawa S, Parhar IS (2015) Role of serotonin in fish reproduction. Front Neurosci 9: 195. doi:10.3389/fnins.2015.00195.

31. Cerdà J, Petrino TR, Greenberg MJ, Wallace RA (1997) Pharmacology of the serotonergic inhibition of steroid-induced reinitiation of oocyte meiosis in the teleost Fundulus heteroclitus. Mol Reprod Dev 48: 282–291. doi:10.1002/(SICI)1098-2795(199710)48:2<282::AID-MRD17>3.0.CO;2-#.

32. Cerdà J, Reich G, Wallace RA, Selman K (1998) Serotonin inhibition of steroid-induced meiotic maturation in the teleost Fundulus heteroclitus: role of cyclic AMP and protein kinases. Mol Reprod Dev 49: 333–341. doi:10.1002/(SICI)1098-2795(199803)49:3<333::AID-MRD14>3.0.CO;2-X.

33. Cerdà J, Subhedar N, Reich G, Wallace RA, Selman K (1998) Oocyte sensitivity to serotonergic regulation during the follicular cycle of the teleost Fundulus heteroclitus. Biol Reprod 59: 53–61.

34. Cerdá J, Petrino TR, Lin Y-WP, Wallace RA (1995) Inhibition ofFundulus heteroclitus oocyte maturation in vitro by serotonin (5-hydroxytryptamine). J Exp Zool 273: 224–233. doi:10.1002/jez.1402730307.

35. Chattoraj A, Seth M, Maitra SK (2008) Influence of serotonin on the action of melatonin in MIH-induced meiotic resumption in the oocytes of carp Catla catla. Comp Biochem Physiol Part A, Mol Integr Physiol 150: 301–306. doi:10.1016/j.cbpa.2008.03.014.

36. Godwin J (2010) Neuroendocrinology of sexual plasticity in teleost fishes. Front Neuroendocrinol 31: 203–216. doi:10.1016/j.yfrne.2010.02.002.

37. Larson ET, Norris DO, Gordon Grau E, Summers CH (2003) Monoamines stimulate sex reversal in the saddleback wrasse. Gen Comp Endocrinol 130: 289–298.

38. Ivashkin E, Khabarova MY, Melnikova V, Nezlin LP, Kharchenko O, et al. (2015) Serotonin mediates maternal effects and directs developmental and behavioral changes in the progeny of snails. Cell Rep 12: 1144–1158. doi:10.1016/j.celrep.2015.07.022.

39. Yang H, Zhou Y, Gu J, Xie S, Xu Y, et al. (2013) Deep mRNA sequencing analysis to capture the transcriptome landscape of zebrafish embryos and larvae. PLoS One 8: e64058. doi:10.1371/journal.pone.0064058.

40. Yan L, Yang M, Guo H, Yang L, Wu J, et al. (2013) Single-cell RNA-Seq profiling of human preimplantation embryos and embryonic stem cells. Nat Struct Mol Biol 20:1131–1139. doi:10.1038/nsmb.2660.

41. Maximino C, P. Costa B, G. Lima M (2016) A Review of Monoaminergic Neuropsychopharmacology in Zebrafish, 6 Years Later: Towards Paradoxes and their Solution. Curr Psychopharmacol 5: 96–138. doi:10.2174/2211556005666160527105104.

42. Yokogawa T, Hannan MC, Burgess HA (2012) The dorsal raphe modulates sensory responsiveness during arousal in zebrafish. J Neurosci 32: 15205–15215. doi:10.1523/JNEUROSCI.1019-12.2012.

43. Prykhozhij SV, Marsico A, Meijsing SH (2013) Zebrafish Expression Ontology of Gene Sets (ZEOGS): a tool to analyze enrichment of zebrafish anatomical terms in large gene sets. Zebrafish 10: 303–315. doi:10.1089/zeb.2012.0865.

44. Croft D, Mundo AF, Haw R, Milacic M, Weiser J, et al. (2014) The Reactome pathway knowledgebase. Nucleic Acids Res 42: D472–7. doi:10.1093/nar/gkt1102.

45. Kairov U, Karpenyuk T, Ramanculov E, Zinovyev A (2012) Network analysis of gene lists for finding reproducible prognostic breast cancer gene signatures. Bioinformation 8: 773–776. doi: 10.6026/97320630008773.

46. Peri S, Navarro JD, Kristiansen TZ, Amanchy R, Surendranath V, et al. (2004) Human protein reference database as a discovery resource for proteomics. Nucleic Acids Res 32: D497–501. doi:10.1093/nar/gkh070.

47. Shih D-F, Hsiao C-D, Min M-Y, Lai W-S, Yang C-W, et al. (2013) Aromatic L-amino acid decarboxylase (AADC) is crucial for brain development and motor functions. PLoS One 8: e71741. doi:10.1371/journal.pone.0071741.

48. Levin M, Buznikov GA, Lauder JM (2006) Of minds and embryos: left-right asymmetry and the serotonergic controls of pre-neural morphogenesis. Dev Neurosci 28: 171–185. doi:10.1159/000091915.

49. Emanuelsson H (1992) Autoradiographic localization in polychaete embryos of tritiated mesulergine, a selective antagonist of serotonin receptors that inhibits early polychaete development. Int J Dev Biol 36: 293–302.

50. Pennati R, Groppelli S, Sotgia C, Candiani S, Pestarino M, et al. (2001) Serotonin localization in Phallusia mammillata larvae and effects of 5-HT antagonists during larval development. Dev Growth Differ 43: 647–656.

51. Hämäläinen M, Kohonen J (1989) Studies on the effect of monoamine antagonists on the morphogenesis of the newt. Int J Dev Biol 33: 157–163.

52. Khozhaĭ LI, Puchkov VF, Otellin VA (1995) [The effect of a serotonin deficiency on mammalian embryonic development]. Ontogenez 26: 350–355.

53. Markova LN, Sadykova KA, Sakharova NI (1990) [The effect of biogenic monoamine antagonists on the development of preimplantation mouse embryos cultured in vitro]. Zh Evol Biokhim Fiziol 26: 726–732.

54. Brustein E, Chong M, Holmqvist B, Drapeau P (2003) Serotonin patterns locomotor network activity in the developing zebrafish by modulating quiescent periods. J Neurobiol 57: 303–322. doi: 10.1002/neu.10292.

55. Sallinen V, Sundvik M, Reenilä I, Peitsaro N, Khrustalyov D, et al. (2009) Hyperserotonergic phenotype after monoamine oxidase inhibition in larval zebrafish. J Neurochem 109: 403–415. doi:10.1111/j.1471-4159.2009.05986.x.

56. Airhart MJ, Lee DH, Wilson TD, Miller BE, Miller MN, et al. (2007) Movement disorders and neurochemical changes in zebrafish larvae after bath exposure to fluoxetine (PROZAC). Neurotoxicol Teratol 29: 652–664. doi:10.1016/j.ntt.2007.07.005.

57. Kawashima T, Zwart MF, Yang C-T, Mensh BD, Ahrens MB (2016) The Serotonergic System Tracks the Outcomes of Actions to Mediate Short-Term Motor Learning. Cell 167: 933–946.e20. doi:10.1016/j.cell.2016.09.055.

58. Pantoja C, Hoagland A, Carroll EC, Karalis V, Conner A, et al. (2016) Neuromodulatory regulation of behavioral individuality in zebrafish. Neuron 91: 587–601. doi:10.1016/j.neuron.2016.06.016.

59. Kim D-K, Tolliver TJ, Huang S-J, Martin BJ, Andrews AM, et al. (2005) Altered serotonin synthesis, turnover and dynamic regulation in multiple brain regions of mice lacking the serotonin transporter. Neuropharmacology 49: 798–810. doi:10.1016/j.neuropharm.2005.08.010.

60. Hasegawa H, Nakamura K (2010) Tryptophan hydroxylase and serotonin synthesis regulation. Handbook of the behavioral neurobiology of serotonin. Handbook of behavioral neuroscience. Elsevier, Vol. 21. pp. 183–202. doi:10.1016/S1569-7339(10)70078-3.

61. Neckers LM, Neff NH, Wyatt RJ (1979) Increased serotonin turnover in corpus striatum following an injection of kainic acid: Evidence for neuronal feedback regulation of synthesis. Naunyn Schmiedebergs Arch Pharmacol 306: 173–177. doi:10.1007/BF00498988.

62. Westerfield M (n.d.) The zebrafish book: a guide for the laboratory use of zebrafish. ci.nii.ac.jp.

63. Spulber S, Kilian P, Wan Ibrahim WN, Onishchenko N, Ulhaq M, et al. (2014) PFOS induces behavioral alterations, including spontaneous hyperactivity that is corrected by dexamfetamine in zebrafish larvae. PLoS One 9: e94227. doi:10.1371/journal.pone.0094227.

64. Kehr J, Yoshitake T (2006) Monitoring brain chemical signals bymicrodialysis. Encyclopedia of Sensors. USA: American ScientificPublishers, Vol. 6. pp. 287–312.

65. Sacchetti P, Sousa KM, Hall AC, Liste I, Steffensen KR, et al. (2009) Liver X receptors and oxysterols promote ventral midbrain neurogenesis in vivo and in human embryonic stem cells. Cell Stem Cell 5: 409–419. doi:10.1016/j.stem.2009.08.019.

66. Subramanian A, Tamayo P, Mootha VK, Mukherjee S, Ebert BL, et al. (2005) Gene set enrichment analysis: a knowledge-based approach for interpreting genome-wide expression profiles. Proc Natl Acad Sci USA 102: 15545–15550. doi:10.1073/pnas.0506580102.

